# Phase-encoded fMRI tracks down brainstorms of natural language processing with sub-second precision

**DOI:** 10.1101/2023.05.29.542546

**Authors:** Victoria Lai Cheng Lei, Teng Ieng Leong, Cheok Teng Leong, Lili Liu, Chi Un Choi, Martin I. Sereno, Defeng Li, Ruey-Song Huang

## Abstract

The human language system interacts with cognitive and sensorimotor regions during natural language processing. However, where, when, and how these processes occur remain unclear. Existing noninvasive subtraction-based neuroimaging techniques cannot simultaneously achieve the spatial and temporal resolutions required to visualize ongoing information flows across the whole brain. Here we have developed phase-encoded designs to fully exploit the temporal information latent in functional magnetic resonance imaging (fMRI) data, as well as overcoming scanner noise and head-motion challenges during overt language tasks. We captured neural information flows as coherent waves traveling over the cortical surface during listening, reciting, and oral cross-language interpreting. The timing, location, direction, and surge of traveling waves, visualized as ‘brainstorms’ on brain ‘weather’ maps, reveal the functional and effective connectivity of the brain in action. These maps uncover the functional neuroanatomy of language perception and production and motivate the construction of finer-grained models of human information processing.

## Introduction

Language plays a central role in human information processing. Existing cognitive models have depicted the flow of information from perception to response, with various processing stages in between (*1–4*). Neural models of language processes suggest that the processing of linguistic information involves a series of steps associated with activations across the brain (*5–7*) via certain processing pathways (*8–10*). One important issue is to determine which processes operate sequentially and which in parallel (*11*). Models describing oral interpreting explicitly suggest there are overlaps between various processes during this complex language task (*12, 13*). Therefore, mapping out the whole-brain dynamics of natural language processing in real time holds the key to the understanding of language and cognitive functioning. Demonstrating how different brain regions interact with each other during language perception and production has been identified as one of the greatest challenges (*14*).

Both spatial and temporal characterizations of brain activities are important for revealing the neural mechanisms of language perception and production and their interaction with domain-general cognitive functions (*15*). Invasive procedures like electrocorticography (ECoG) or implanted electrodes have recorded the neural activities of language processing with high spatial and temporal resolution from local regions, but cannot record whole-brain activity (*16–20*). Existing noninvasive neuroimaging techniques such as magnetoencephalography (MEG) and functional magnetic resonance imaging (fMRI) can only achieve either millisecond or millimeter precision, but not both (*7, 14, 21*). Due to its lower temporal resolution and the technical challenges, including scanner noise and speaking induced-head motion artifacts (*22*), fMRI has been somewhat overlooked for capturing the continuous processing of natural language comprehension and production at the second-to-second level. Sparse-sampling fMRI can localize brain activations associated with overt vocal responses during silent delays between acquisitions (*23*), but cannot reveal the temporal order of the ongoing brain activities within and beyond these regions.

Several studies have used time-resolved fMRI to determine the activation sequences of distinct brain regions during language perception and production or bilingual control (*24–27*). However, how information flows continuously from region to region across the language network remains less clear. Also, there is still a lack of direct evidence for the understanding of feedback at the speech production stage. In this study, we used real-time phase-encoded fMRI to capture and unravel the spatiotemporal traveling waves of blood-oxygen-level-dependent (BOLD) signals across the cortical surface during naturalistic language tasks, which involved overt speech. Phase-encoded designs have long been used to map the topological organization of human occipital, parietal, frontal, auditory, and sensorimotor cortices (*28–33*). In a typical retinotopic mapping experiment, for example, a wedge containing flickering checkerboard pattern or live-feed videos slowly revolves around a central fixation for a period of 64 s (*29*, *31*, *32*). Cortical patches representing different polar angles are thus stimulated at different phases. Fourier analysis of resulting fMRI time series reveals periodic brain activities at the stimulus frequency (e.g., 8 cycles/scan), often visualized as surface waves traveling across polar-angle representations within each retinotopic area (*29, 31, 32*). The very slow rate of stimulus progression was initially chosen to respect the slow hemodynamic response. However, it was yet to be determined whether phase-encoded designs could be used to capture faster brain dynamics during real-time language processing. A recent phase-encoded fMRI study demonstrates that the activation sequence of sensorimotor transformations during a periodic reach-to-eat task could be decoded within a period as short as 16 s (*34*). Inspired by this recent success, we conducted an fMRI experiment with a series of rapid phase-encoded tasks involving unimodal (reading or listening in Chinese [native language, L1] or English [second language, L2]), multimodal (reading aloud or shadowing in L1 or L2), crossmodal (reading or listening and then reciting in L1 or L2), and cross-language (crossmodal L1-to-L2 or L2-to-L1) processing of natural sentences (Figs. 1, 2, Fig. S1). Twenty-one native Chinese speakers with English as their second language participated in this fMRI study. Each 256-s functional scan consisted of 16 periodic trials, where a written or spoken sentence (Tables S1, S2) was presented between 0 and 5 s within each 16-s trial. The subjects followed visual cues to read, listen, read aloud, or shadow each sentence (Fig. 1A to D). In crossmodal and cross-language scans, subjects were required to memorize a written or spoken sentence, or a sequence of five digits, and then recite or orally translate it between 5-10 s within each trial (Figs. 1E, F, 2). The subjects were able to clearly hear their own speech in real time via active noise-canceling headphones. To ensure task compliance, the subject’s voice and eye movements were recorded continuously during scanning using MR-compatible devices (Figs. S2, S3, S4, Table S3). To prevent head motion, especially in tasks involving speaking, subjects wore “The Phantom of the Opera” style masks molded to individual faces throughout each fMRI session (Figs. S2, S5).

**Figure 1.**
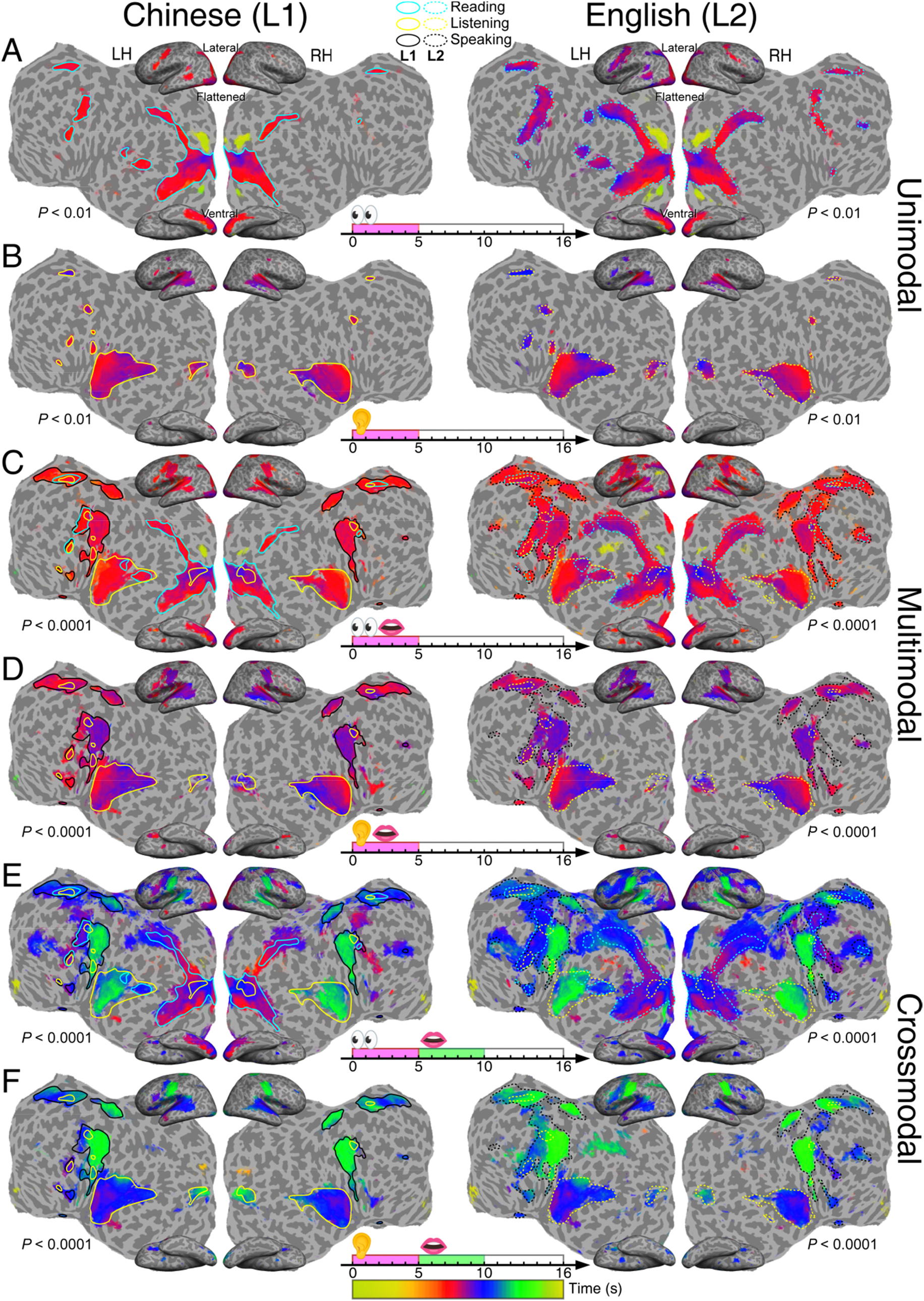
Group-average maps (n = 21) of periodic activations in single-language tasks. (**A**) Silent reading (0-5 s). (**B**) Listening (0-5 s). (**C**) Reading aloud (0-5 s). (**D**) Shadowing (listening to and immediately repeating a speech within 0-5 s). (**E**) Reading and memorizing a written sentence (0-5 s) then reciting it (5-10 s). (**F**) Listening to and memorizing a spoken sentence (0-5 s) then reciting it (5-10 s). Significant periodic activations are displayed on the inflated (lateral and ventral) and flattened cortical surfaces of a representative subject. L1: Chinese; L2: English. LH: left hemisphere; RH: left hemisphere. The colorbar indicates the phase angles (0-2π) of BOLD signals corresponding to a trial period (0-16 s). A lower statistical significance threshold (*F*_(2, 230)_ > 4.7; equivalent *P* < 0.01, uncorrected) is set for the maps of unimodal (reading and listening) tasks to match the spatial extent of significant activations (*F*_(2, 230)_ > 9.6; equivalent *P* < 0.0001, uncorrected) in multimodal and crossmodal tasks. Color contours indicate regions activated by silent reading (cyan), listening only (yellow), and speaking (black). Solid contours: L1 activations; dotted contours: L2 activations.

**Figure 2.**
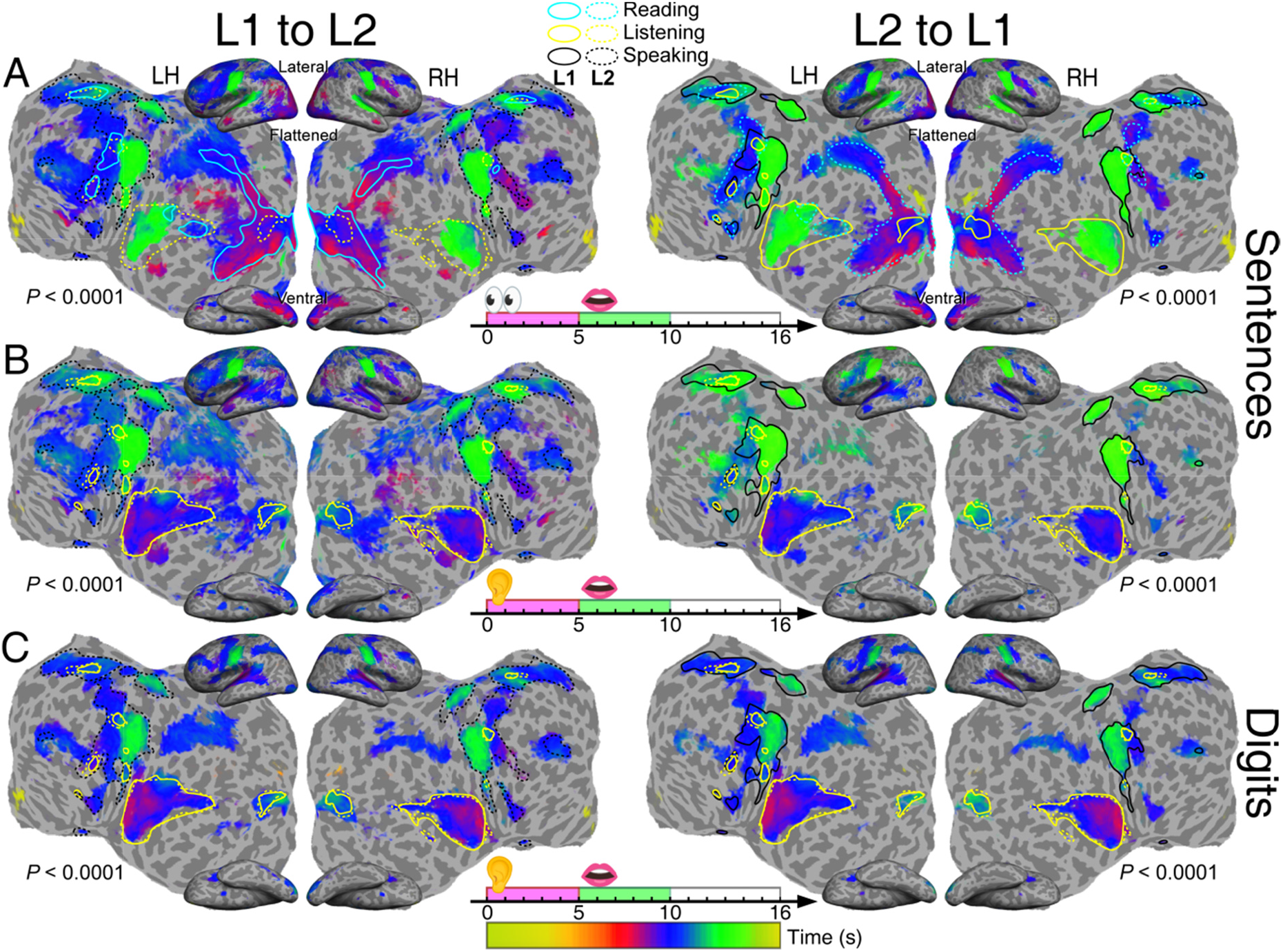
Group-average maps (n = 21) of periodic activations in translation tasks. (**A**) Text-to-speech translation (reading and memorizing a written sentence in 0-5 s, then verbalizing it in the target language in 5-10 s). (**B**) Speech-to-speech translation (listening to and memorizing a sentence in 0-5 s, then vocalizing it in the target language in 5-10 s). (**C**) Digit interpreting (listening to and memorizing five digits in 0-5 s, then vocalizing them in the target language in 5-10 s). All maps are displayed with a statistical significance *F*_(2, 230)_ > 9.6 (equivalent *P* < 0.0001, uncorrected). Color contours and all other conventions are the same as those from Fig. 1.

## Results

### Dynamic brain ‘weather’ maps

Time series of fMRI data were analyzed with Fourier-based and surface-based averaging methods (*29–34*) (see **Materials and Methods**). Group-average maps of statistically significant periodic activations at 16 cycles/scan and their phases are displayed in an orange-red-blue-green-yellow phase gradient on the flattened cortical surfaces of a representative subject (Figs. 1, 2). The spatiotemporal patterns of surface traveling waves in four representative tasks are illustrated as isophase contour maps (Fig. 3). To interpret the traveling wave patterns, we used the topological maps recently published by Sereno et al. as a reference map for locating and labeling functional brain areas (*33*, Fig. 4A). For easy comparison of activated areas between different tasks and languages, we overlaid the contours of L1 and L2 modality-specific activations on the topological maps and summarized the directionality of traveling waves in Fig. 4B, C. The visualization of spatiotemporal unfolding of traveling wave patterns on brain ‘weather’ maps (Figs. 3, 4) and ‘brainstorm’ movies (Movies S1 to S18) is analogous to the tracking of dynamic rainbands across state and county lines on a dynamic weather radar map. The color-coded maps here represent surface traveling waves (*29–33*) propagating within and across fixed cortical area borders (cf. county lines) of topological and non-topological areas during real-time language processing. Arrows indicate the local direction of the coherent traveling waves in static pictures (Figs. 3E to P, 4B, C).

**Figure 3.**
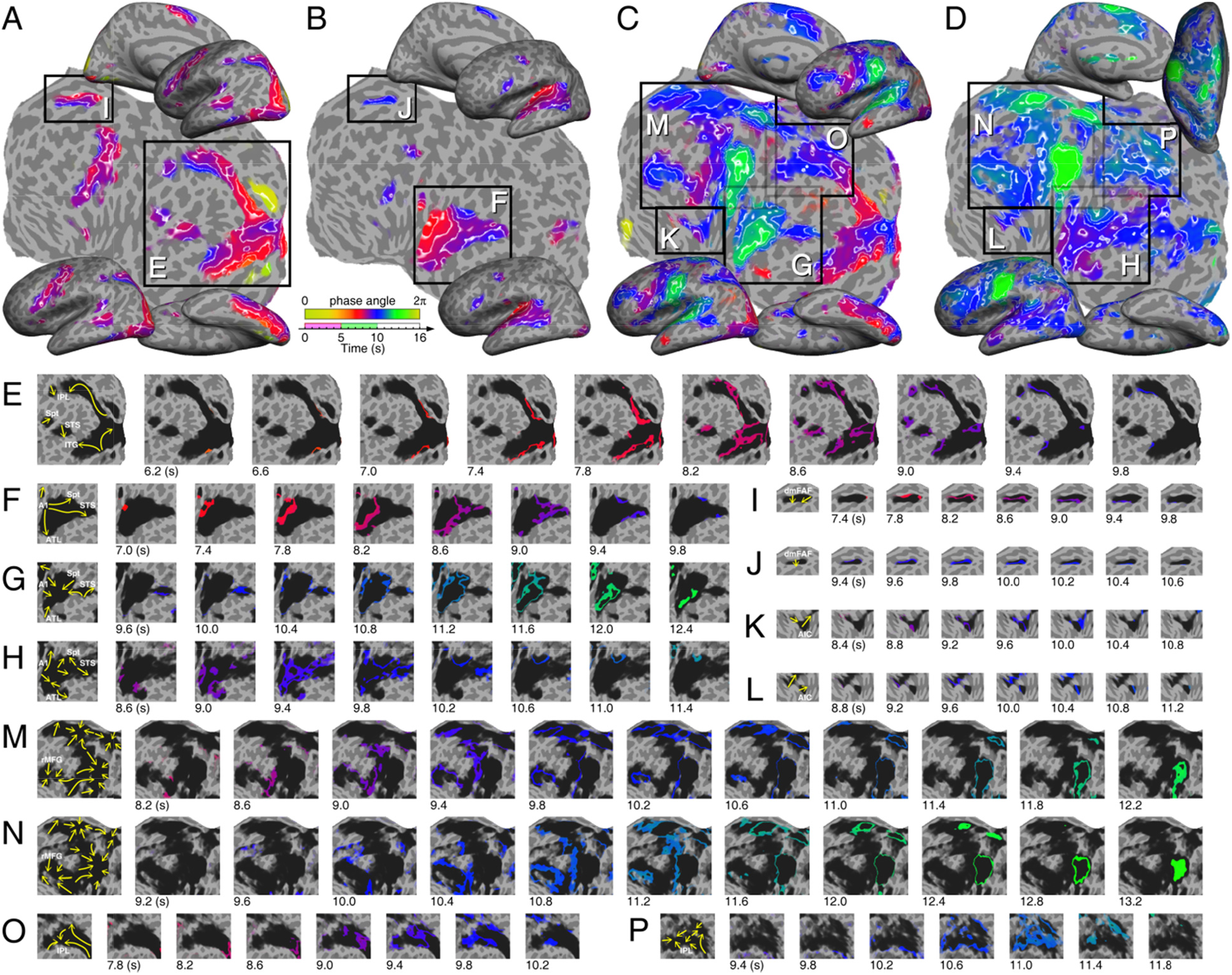
Tracking the paths of traveling waves on isophase contour maps. (**A**) L2 reading task (Fig. 1A, right). (**B**) L2 listening task (Fig. 1B, right). (**C**) L1 reading-memorizing-r**e**citing task (Fig. 1E, left). (**D**) L1 (listening) to L2 (verbalizing) interpreting task (Fig. 2B, l**e**ft). (**E-P**) Each frame sequence shows the traveling waves (cf. storm rainbands) in a corresponding box in (A-D). The isophase of traveling waves in each frame is color-coded and displayed over dark ‘clouds’ representing the extent of activations in (A-D). Arrows in the leftmost frame of each sequence represent the paths of local traveling waves.

**Figure 4.**
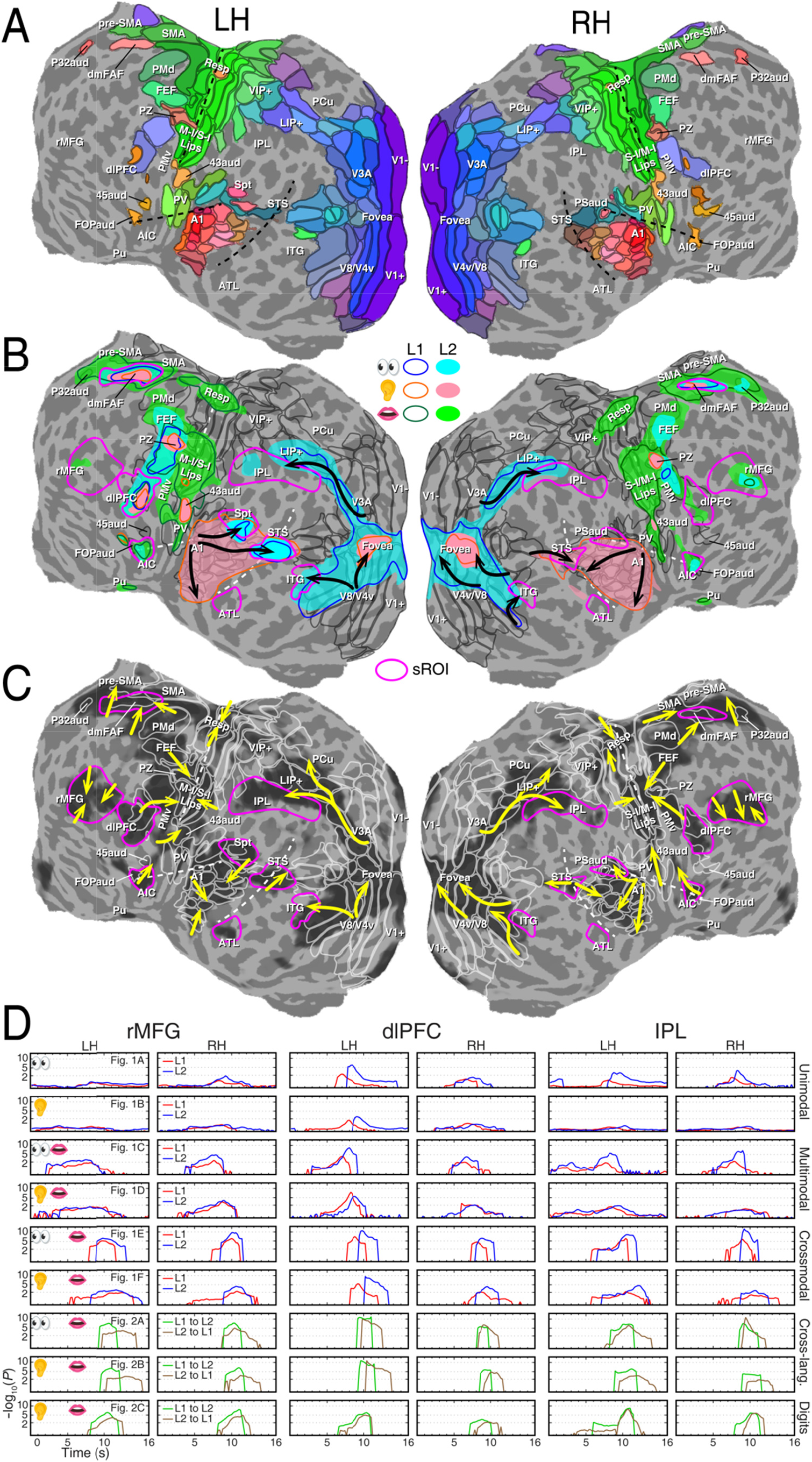
Tracking traveling waves and surges on multilayer brain ‘weather’ maps. (**A**) Topological maps with dark gray contours indicating the borders of visual, auditory and somatomotor areas (*33*) displayed on the flattened cortical surfaces of a representative subject (see Table S4 for full names of labels). (**B**) Multilayer single-language activation maps overlaid on topological areas. Color contours (L1) and filled regions (L2) outline the extent of unimodal (reading and listening) or multimodal (speaking and self- monitoring) activations in Fig. 1. Black arrows indicate multiple streams of traveling waves in visual and auditory cortices. (**C**) Tracking the paths (yellow arrows) of regional traveling waves over topological areas (white contours) during a L1 reading-memorizing-reciting task (Fig. 1E, left). Dark ‘clouds’ represent the extent of significant periodic activations (*F*_(2, 230)_ > 7.1; equivalent *P* < 0.001, uncorrected). Magenta contours indicate sROIs of language and domain-general cognitive regions outside or partially overlapping topological areas. (**D**) Surge profiles of traveling waves arriving and leaving six bilateral sROIs (rMF, dlPFC, and IPL). Each row corresponds to a task in Figs. 1 and 2. The height of surge is indicated by -log_10_(uncorrected *P*-value) in the y-axis. Red and blue profiles indicate L1 and L2 single- language tasks respectively. Green and brown profiles indicate L1-to-L2 and L2-to-L1 translation tasks respectively.

### Visualizing surface traveling waves

Figure 1 shows the phase-encoded activation maps of unimodal (involving a single sensory modality), multimodal (involving two or more sensory and motor modalities concurrently), and crossmodal (changing modalities) tasks in Chinese or English. Multimodal and crossmodal tasks always elicited more extensive activations than unimodal tasks. Figure 2 shows the phase-encoded activation maps for six translation tasks. For sentence translation, the L1-to-L2 direction (Fig. 2A, B) always induced slightly larger activations than the L2-to-L1 direction. Similarly, for digit translation, the L1-to- L2 direction (Fig. 2C) induced a slight increase in activated areas compared with the L2-to-L1 direction.

To assist interpretation of the intrinsic spatiotemporal brain dynamics during unimodal, crossmodal, and cross-language tasks, we produced a series of ‘brainstorm’ movies (Movies S1 to S18) illustrating the paths of traveling waves across flattened cortical surfaces. For example, in the L1 reading- memorizing-reciting task (Figs. 1E, 3C; Movie S5), waves of activations first emerged in early visual areas and the anterior temporal lobe (ATL) during reading. The occipital waves then branched off into dorsal and ventral visual streams, reaching up to lateral intraparietal areas (LIP+) and the inferior temporal gyrus (ITG), respectively. Around the time when the waves reached LIP+ and ITG, the premotor cortex, anterior insular cortex (AIC), dorsolateral prefrontal cortex (dlPFC), and rostral middle frontal gyrus (rMFG) were activated. As the waves from the premotor cortex and AIC continued to travel toward oral sensorimotor areas, the pre-supplementary motor area (pre-SMA) was also activated. The waves then propagated from pre-SMA to SMA in preparation for speech output. Lastly, as the subjects started vocalizing, the sensorimotor and auditory cortices were activated simultaneously, reflecting self-monitoring during speech production.

The activation patterns in each movie corresponding to a language task (Figs. 1, 2) are described in detail below. In reading tasks (Fig. 1A; Movies S1, S7), activation waves emerged from early visual areas, and branched off into the dorsal and ventral visual streams (*35*) reaching up to LIP+ and ITG (Figs. 3A, E, 4B). Smaller clusters of activations were also present in the superior temporal sulcus (STS), Sylvian parietal temporal area (Spt) (*36*), an area overlapping both pre-SMA and SMA (labeled separately as dorsomedial frontal auditory field, dmFAF) (*33*), dlPFC, frontal eye fields (FEF), and premotor cortex. In listening tasks (Fig. 1B; Movies S2, S8), activation waves emerged from the primary auditory cortex (A1), and branched off into three streams (*8, 9*) reaching: (1) superior temporal gyrus (STG) and Spt, (2) STS, and (3) ATL (Figs. 3B, F, 4B). Other smaller clusters of activations were observed in pars triangularis, dlPFC, auditory polysensory zone (PZ), area 43aud, and SMA (Fig. 4B). On the whole, the activations were predominantly bilateral for both English and Chinese reading and listening tasks.

Multimodal reading-aloud tasks (Fig. 1C; Movies S3, S9) involved reading, speaking, and self-monitoring of speech production simultaneously. Correspondingly, additional activations as compared with reading (Fig. 1A; Movies S1, S7) and listening tasks (Fig. 1B; Movies S2, S8) were found in SMA, primary and secondary sensorimotor cortex (oral and diaphragm representations), pars triangularis, STG, and AIC (outlined by black contours in Fig. 1C). Multimodal shadowing tasks (Fig. 1D; Movies S4, S10) involved listening to stimuli, speaking, and self-monitoring simultaneously. Additional activations were found in pre-SMA, sensorimotor cortex, and AIC as compared with listening tasks (Fig. 1B). Compared with reading-aloud tasks (Fig. 1C), shadowing tasks activated broader areas in the STS bilaterally. Although the subjects were instructed to speak as soon as the stimuli appeared, delays in activation (purplish) were found around the premotor and sensorimotor areas in shadowing tasks (Fig. 1D) for both English and Chinese. The audio recordings confirmed that shadowing tasks induced delays in speech onset and offset compared with reading-aloud tasks (Table S3). Similar to the unimodal maps, activations were predominately bilateral and the L2 maps show slightly increased activated areas compared with the L1 maps (Fig. 1D).

Crossmodal maps (Fig. 1E, F) show sequential activations before, during, and after modality- change, indicating that the subjects had to first read silently or listen to the English or Chinese sentences, commit them into memory, and then recite them. In reading-memorizing-reciting tasks (Figs. 1E, 3C; Movies S5, S11), the three modalities (reading, speaking, and listening to own speech) each sequentially activated specific brain regions as revealed by the traveling waves, with the early visual areas being first activated (reddish regions). Purple-bluish regions indicate intermediate activations, which were mainly involved during the modality-switch (between reading and speaking). Additional activated areas as compared with reading-aloud tasks (Fig. 1C) include ATL, rMFG, and the inferior parietal lobe (IPL) (Figs. 3C, M, O, 4C). Lastly, greenish regions indicate late activations (Fig. 3C, G, M), which were mainly clustered in the primary sensorimotor areas (oral and diaphragm), auditory cortex, and Spt as the tasks involved speech motor control and self-monitoring. The smaller extent of activation in the auditory cortex suggests its involvement in hearing and monitoring of self-speech (Fig. 1E; yellow outline). Similar to the unimodal and multimodal maps, the crossmodal activations were bilateral. Compared to L1 maps, L2 maps overall show significantly larger extent in activation for the reading-memorizing-reciting tasks (Fig. 1E; right panel). On the other hand, during the listening-memorizing-reciting tasks (Fig. 1F; Movies S6, S12), A1 and ATL were the first areas to be activated. Intermediate activations included higher-level auditory cortex, AIC, premotor, and pre-SMA regions. Late activations included fovea, IPL, Spt, and the primary sensorimotor areas. The L1 and L2 maps demonstrate similar patterns except for the activations in the premotor and IPL regions, where activations induced by L2 tasks are larger than those associated with L1 tasks.

Figure 2 shows the spatiotemporal patterns of cross-language control in translation tasks. The maps were organized by stimulus type (sentences or digit sequences) and translation direction (L1-to-L2 or L2-to-L1). Color contours were taken from the activations found in reading only, listening only, and speaking (reading aloud) tasks (Fig. 1A, B, C). All activation patterns in the reading-memorizing-reciting tasks were found in the reading-memorizing-translating tasks (Fig. 2A, Movies S13, S16). Similarly, activations in the listening-memorizing-reciting tasks were also present in the listening-memorizing- translating tasks (Fig. 2B, Movies S14, S17).

In sentence-level translation tasks, the L1-to-L2 direction involved extensive activations in the posterior middle temporal gyrus (MTG), posterior STS, IPL, dlPFC, rMFG, and superior frontal gyrus. On the other hand, L2-to-L1 translation tasks exhibited smaller activation extent in domain-general cognitive regions and very sparse activations in the posterior MTG and STS. In addition, when compared to the single-language crossmodal tasks, the cross-language processing exhibited more extensive activations in ATL, posterior MTG/STS, and the domain-general cognitive regions, including IPL and prefrontal regions. In digit-level translation tasks, the L1-to-L2 direction slightly expanded the surface area of the activated regions compared with L2-to-L1 digit translation (Fig. 2C, Movies S15, S18). In addition, the extent of activations in the STS was similar to those during reading-aloud and reading- memorizing-reciting tasks (Fig. 1C, E). Besides a smaller activation in STS, ATL and posterior MTG/STS were not activated in digit-level translation tasks (Fig. 2C) as compared with the verbal sentence-level translation tasks (Fig. 2B), which may help confirm the roles of STS, ATL and posterior MTG/STS in semantic processing and verbal reasoning. Lastly, activations were found in IPL, rMFG, and dlPFC, suggesting the involvement of working memory in translation.

### Analyzing local traveling wave patterns

One goal of this study is to present a new way to analyze and visualize spatiotemporal interactions within and across the sensory, motor, language, and cognitive brain regions. We selected four representative tasks to illustrate the complex spatiotemporal patterns of local traveling waves (Fig. 3A to D as Movies S7, S8, S5 and S14 respectively). Figure 3 (A, E) shows three streams of waves in the ventral and dorsal visual pathways during an L2 reading task (Figs. 3A, 4B). Two waves emerged from areas V8 and V4v, with one propagating to ITG and the other to foveal representations in early visual areas. The third wave emerged from area V3A and traveled upstream along the LIP+ cluster to reach IPL. All tasks that involved reading triggered and displayed similar patterns and directions of traveling waves in ventral and dorsal visual pathways (Figs. 1A, C, E, 2A). Figure 3 (B, F) shows that activation waves emerged from A1 and split into at least three streams reaching ATL, Spt, and STS in the left hemisphere during an L2 listening task (Fig. 4B). Interestingly, during the production of speech from visually presented sentences such as the reading-memorizing-reciting tasks (Figs. 1E, 2A) and reading- memorizing-translating tasks (Fig. 2B), the direction of waves emerging from A1 remained the same, while the other two streams showed reverse directions, i.e., they initiated from Spt and STS and propagated anteriorly (compare Fig. 3F, G).

Regarding domain-general cognitive regions, traveling waves initiated from the perimeter of rMFG and propagated radially inwards (Fig. 3M, N). Similar radial-inward flow patterns were found in IPL (Fig. 3O, P). These patterns remained identical across all the tasks that recruited rMFG and IPL. Similarly, the patterns in AIC and dlPFC remained identical for all the tasks that recruited these regions (Fig. 3K, L). Most remarkably, an alignment of planar wavefronts emerged from AIC and dlPFC, which then merged at the border between premotor and sensorimotor areas (Fig. 3K to N). Furthermore, distinct spatiotemporal wave patterns were observed in dmFAF: one downward stream was found in the unimodal tasks (Fig. 3I, J); two radial-inward streams that each emerged from the perimeter and propagated towards the center were observed in the multimodal and crossmodal tasks (Fig. 3C, D, M, N).

### Regional surges of traveling waves

Once the paths of traveling waves were identified, we then selected nine surface-based regions of interests (sROIs) in domain-general cognitive regions and language-related regions, including rMFG, dlPFC, IPL, dmFAF, AIC, Spt, ATL, STS, and ITG (magenta contours in Fig. 4B, C), to compare their involvement in the unimodal, multimodal, crossmodal, and cross-language tasks. Figure 4D and Fig. S6 show the distribution of phase-sorted complex F-values within each sROI, which reveals the surge, subside, latency, and peak magnitude of the traveling waves (cf. regional storm surges) of corresponding tasks in Figs. 1 and 2. For example, dlPFC was predominantly activated in the left hemisphere across all tasks. The highest activation surges occurred during the crossmodal and sentence translation tasks. Three different latencies were found across all tasks. For the unimodal tasks and digit translation tasks, there were medium activation delays. The delays were short for both of the multimodal tasks. The crossmodal and sentence translation tasks showed the longest delay. L1 production exhibited shorter latency than L2 production while L1-to-L2 translation exhibited slightly shorter latency than L2-to-L1 translation.

## Discussion

Temporal and spatial measures must be combined to acquire a deeper understanding of the neural basis of language processing. Previous time-resolved fMRI studies have revealed sequential activations of multiple isolated brain regions, but without showing continuous spatial and temporal progression across regions (*24–27*). How activation arises in one brain region and propagates towards the next region during real-time language processing has remained obscure. In visual neuroscience, phase-encoded fMRI mapping using time-delayed retinotopic stimuli reveals spatially and temporally continuous traveling waves across the cortical surface (*28–33, 37*). However, the rate of stimulation (e.g., a period of 64-s) in typical phase-encoded designs is too slow for mapping real-time language perception and production, where a sentence is usually processed within a few seconds. In this study, we developed rapid phase-encoded fMRI to capture and visualize fine-grained spatiotemporal BOLD signals propagating over the cortical surface during real-time naturalistic unimodal, crossmodal, and cross-language processing. We demonstrated that 5 s of naturalistic reading or listening induced coherent surface waves that traveled continuously across the visual or auditory streams, which had not been revealed in previous time-resolved fMRI studies of higher cognitive processes (*24–27, 38*). The tracking of traveling waves in ‘brainstorm’ movies, showing the brain in action, opens a new window into the understanding of the dynamic functional and effective connectivity across the whole brain during real-time language processing. The timing, location, and direction of traveling waves reveal information flow within and across visual, auditory, sensorimotor, language, and cognitive regions, which can be integrated into existing models of human information processing. Major findings of our study are discussed below.

Across all the experimental tasks for this study (Figs. 1A to F 2A to C), we identified a bilateral network of task-induced activations during natural language processing, with certain tasks such as consecutive interpretation of sentences exhibited a slight bias to the left hemisphere. It is a long standing view that language is predominantly left lateralized, though it has been suggested that the right hemisphere also has a role to play in language processing. Findings from functional neuroimaging studies regarding language lateralization are contradictory. This study gives the most direct and consistent support to the recently proposed bilateral model of language processing, which suggests that the language network is asymmetric but intrinsically bilateral (*39*).

In Figs. 1A and 3E, the parallel traveling waves across the ventral and dorsal visual cortex suggest that reading involves text localization (where) as well as recognition (what), which provides direct evidence for the dual-stream hypothesis of visual processing(*35*). Furthermore, the spatiotemporal patterns in the two visual pathways were consistent across all tasks involving reading (Figs. 1A, C, E, 2A). During a passive listening task, traveling waves emerged from A1 and split into roughly three streams reaching Spt, STS, and ATL in the left hemisphere (Figs. 1B, 3F, 4B). The spatiotemporal patterns in the auditory cortex were similar for all the tasks involving listening to externally delivered speeches (Figs. 1B, D, F, 2B, C). The flow patterns within the auditory cortex also provide direct evidence for, at minimum, a dual-stream model of auditory speech processing (*8, 9*).

During the reading-aloud task (Fig. 1C), the flow patterns in the visual cortex remain consistent with those during silent reading tasks (Fig. 1A). Similarly, in the shadowing tasks (Fig. 1D), we observed three streams of waves emerging from A1. The consistent and reproducible traveling wave patterns are likely attributable to the underlying functional dynamics (*40*). Compared with unimodal tasks, the extended activations and distinct flow patterns in SMA during simultaneous multimodal tasks (Fig. 1C, D) reflect additional recruitment for speech motor preparation (*41*). Activations with the same phases (green) in premotor and sensorimotor cortices and Spt also reflect motor control and self-monitoring of speech production (*36, 42, 43*).

In the tasks involving both speech perception and production (Fig. 1D, F), the phases of activations in the superior temporal cortex were dominated by external stimulation and very similar to those in the passive listening tasks (Fig. 1B). However, different flow patterns of activations in higher-level auditory cortex originating from STS were observed during self-monitoring of speech in tasks involving reading aloud or reciting (Figs. 1C, E, 2A). This suggests that reading for speech production and self-monitoring engages different neural mechanisms than the perception of external speech.

During the simultaneous multimodal tasks, the activation phases of the visual or auditory cortex were less distinguishable from those of the oral sensorimotor cortex (Fig. 1C, D). One of the advantages of phase-encoded designs over non-time-resolved block designs or event-related designs is that the delayed reciting in crossmodal tasks not only separates the activation phases of language input (reading or listening) from that of speech output, but also recruits domain-general cognitive regions involved in working memory and other executive functions (Fig. 1E, F). Interestingly, the activation maps for crossmodal tasks seem to indicate that reading-memorizing-reciting tasks involve more cognitive effort than listening-memorizing-reciting tasks. This reflects functionally distinct mechanisms for translating written to verbal language and temporary buffering of verbal information before speech output prompted by a visual cue. Furthermore, the additional activations in bilateral domain-general cognitive regions were more prominent in L2 tasks (Fig. 1E, F; right panel), suggesting that processing L2 is more cognitively demanding (see Fig. 4D and discussion below). Lastly, Spt in the left posterior Sylvian fissure displayed phases coherent with the oral and diaphragm sensorimotor areas in the crossmodal tasks (greenish regions in Fig. 1E, F). This suggests that Spt is involved in the processing of linguistic information for speech production and monitoring, supporting its role in sensory-motor integration (*36*).

During a reading-memorizing-reciting task, traveling waves sequentially propagate through the occipital, parietal, temporal, and prefrontal cortices, and finally reach speech sensorimotor and auditory cortices (Figs. 1E, 3C, 4C). These spatiotemporal flow patterns provide empirical evidence for logistics models of information processing in the brain (*33, 44*). Understanding a written sentence may involve ‘serial assembly of content’ operation – similar to scene comprehension from a series of fixations, which mirrors how connected speech is parsed into chunks for analysis in the brain (*44*). Individual words are first ‘packed and carried’ from the early visual cortex into ventral and dorsal visual streams by traveling waves through retinotopic areas (Fig. 4A, C). The assembled information may then ‘escape’ from the topologically mapped cortex and ‘spill over’ into more domain-general cognitive regions (e.g., IPL, dlPFC, and rMFG) involved in working memory and other executive functions, while still exhibiting spatially coherent wave-like activation patterns (*33*). After brief ‘storage-in-transit’ in these regions, the information then flows back into the topologically organized somatomotor cortex to generate speech output, which is simultaneously monitored by the auditory cortex during vocalization. Similar patterns of waves flowing in and out of topologically mapped cortex were observed in the crossmodal and cross-language tasks (Figs. 1, 2). However, a larger extent of ‘overspill’ into domain-general cognitive regions (e.g., IPL) was observed in the cross-language tasks, suggesting that they involved greater cognitive effort.

Moving outside of the topological areas, the surge profile of each sROI provides additional information to compare the arrival, ending, duration, and height of traveling waves within each domain-general cognitive region across different language tasks (Fig. 4D, Fig. S6). In contrast, regions identified by non-time-resolved general linear models only show statistically significant activation strength and extent without resolving the temporal information of activations within and between them. In most single-language tasks, both L1 and L2 exhibited similar activation duration as shown in the surge profiles of the same sROI (e.g., dlPFC). L1 tasks always activated domain-general cognitive regions earlier than L2 tasks, while L2 tasks induced higher magnitudes, suggesting that L2 processing is indeed more cognitively demanding. During the crossmodal cross-language processing, domain-general cognitive regions in L1-to-L2 translation tasks triggered activations earlier than in L2-to-L1 translation tasks, and L2-to-L1 translation tasks induced lower but more extended activation. This suggests that L2-to-L1 processing may require lesser but prolonged cognitive effort. Taken together, phase-encoded fMRI enables the analysis of temporal evolution of activations in domain-general cognitive regions. The surge profiles of traveling waves can serve as potential indicators of cognitive performances in language and non-language tasks.

In this study, we have overcome limitations of existing fMRI techniques, e.g., loud gradient-coil noise and speaking-induced head motion artifacts (*22*), and have conducted real-time fMRI during naturalistic language tasks (reading aloud, listening, shadowing, and consecutive interpreting), unveiling the many facets of human information processing. Maximum efforts were made to enhance ecological validity of the study – besides using natural sentences as stimuli and involving overt speech, we also simulated the real-world situation of consecutive interpreting by giving a visual cue to prompt the subject’s response. Very importantly, we have solved the temporal bottleneck problem of fMRI by using rapid phase-encoded designs and simultaneously visualized the spatial and temporal unfolding of surface traveling waves across the entire brain during real-time unimodal, multimodal, crossmodal, and cross-language tasks. The unfolding of hemodynamic traveling waves, albeit much slower, is strongly reminiscent of neural oscillations (*45, 46*), suggesting the waves originate from neuronal activities. By analyzing the spatiotemporal traveling wave patterns, we can reveal the real-time functional connectivity between brain regions activated at each moment (i.e., local and distant activation pattern in each frame of a ‘brainstorm’ movie) as well as hints of the direction of information flow and possible causality between brain regions (as indicated by arrows in Figs. 3, 4), thus providing direct evidence for multiple human information processing models in visual, auditory, and sensorimotor cortices (*8, 9, 14, 35, 47–49*). We believe that rapid phase-encoded fMRI can be adapted outside language studies. The ability to track down the step-by-step cognitive processes across the whole brain at sub-second precision may advance our understanding of the computational principles of the widely distributed cognitive network, in both normal and abnormal brain functioning.

This study provides a more fine-grained picture of spatiotemporal information flow in the brain during natural language functioning. To share these results, we have constructed an open access database, the University of Macau Brain Atlas (UMBA; Movie S19; see **Materials and Methods**), which incorporates annotated multilayer, multimodal, multi-language, and multi-population functional activation maps and ‘brainstorm’ movies. Similar to interactive weather websites with dynamic radar maps, UMBA shows the directions of traveling waves (cf. storms paths) during real-time language processing across the flattened cortical surface. UMBA will undergo continuous updates with new layers of maps and movies from different languages, tasks, and populations, in the hope that it can be used as a Google Earth-like interactive guide by the neuroimaging community (Movie S20).

## Materials and Methods

### Participants

Twenty-one bilingual students (15 females; 6 males) between 20 and 42 years old (Mean: 23.5; SD: 4.9) participated in this study. All of them were native Chinese speakers with English as their second language, and they started learning English between the age of 3 and 13 (Mean: 7.5; SD: 2.6). Their levels of English proficiency were evaluated based on their recent results in standardized English language tests (IELTS 6.5+ or equivalent). All of them had normal or corrected-to-normal vision and no history of neurological impairment. All subjects gave written informed consent according to experimental protocols approved by the Ethics Committee of the University of Macau.

### Stimuli

Chinese (L1) stimuli included a total of 128 sentences, each consisting of 14 Chinese characters (see examples in Table S1). For each sentence, a subject, verb and an object were selected from *The Lancaster Corpus of Mandarin Chinese* (https://www.lancaster.ac.uk/fass/projects/corpus/LCMC/) to construct simple sentences in the subject-verb-object word order. All sentences were constructed systematically by native Chinese speakers and validated by a professor of linguistics to ensure all sentences were plausible and unambiguous. English (L2) stimuli included a total of 128 sentences, each consisting of 15 to 17 syllables (see examples in Table S2). For the construction of each sentence, a verb was first selected from *Longman Communication 3000,* a list of 3000 most frequently used words in spoken and written English. The verb was then searched in *Collins Dictionary English* (https://www.collinsdictionary.com/dictionary/english/corpus) for sentence examples to ensure the sentences were natural. Only sentence examples that are in the simple subject-verb-object structure were selected and modified to meet the syllable count requirement. Finally, the modified sentence was checked with *Longman Vocabulary Checker* (http://global.longmandictionaries.com/vocabulary_checker) to ensure all the words in the sentence were middle to high frequency words. All sentences were then validated by a professor of linguistics to ensure they were unambiguous and plausible.

### Experimental design and paradigm

Each subject participated in three sessions – one Chinese language session, one English language session, and one translation (L1-to-L2 and L2-to-L1) session. The three sessions were on three different days. Each of the Chinese or English sessions consisted of twelve 256-s functional scans, two scans for each of the six different tasks:

Scan #1: Reading the visually presented sentence silently;

Scan #2: Reading aloud the sentence as soon as it is visually presented;

Scan #3: Reading silently and memorizing the visually presented sentence, then reciting it when a mouth cue appears;

Scan #4: Reading the visually presented sentence silently;

Scan #5: Reading aloud the sentence as soon as it is visually presented;

Scan #6: Reading silently and memorizing the visually presented sentence, then reciting it when a mouth cue appears;

Scan #7: Listening to the auditorily presented sentence;

Scan #8: Shadowing the auditorily presented sentence (repeating what is heard aloud right after the onset of the auditory presentation of the sentence);

Scan #9: Listening to and memorizing the auditorily presented sentence, then reciting it when a mouth cue appears);

Scan #10: Listening to the auditorily presented sentence;

Scan #11: Shadowing (repeating what is heard aloud right after the onset of the auditory presentation of the sentence);

Scan #12: Listening to and memorizing the auditorily presented sentence, then reciting it when a mouth cue appears).

Each 256-s scan consisted of sixteen randomly selected sentences, each sentence was presented visually (Scans #1 to #6) or auditorily (Scans #7 to #12) within the first 5 s of each 16-s trial period (Fig. S1). For Scans #1 to #6, each sentence was presented in black text (32 pt. Arial font), centered against a white background on an LCD monitor. For Scans #1 and #4, each sentence was presented for 5 s with a visual cue (eyes) above the sentence, followed by a blank screen for 11 s within each 16-s period (Fig. S1A). Subjects were instructed to read the sentence silently and attentively within the first 5 s and then rest for 11 s with their eyes open. For Scans #2 and #5, each sentence was presented for 5 s with a visual cue (eyes and mouth) above the sentence followed by a blank screen for 11 s (Fig. S1C). Subjects were instructed to read aloud the sentence as soon as it appeared within 5 s and rest for 11 s with their eyes open. For Scans #3 and #6, each sentence was presented for 5 s with a visual cue (eyes) above the sentence, followed by a visual cue (mouth) for 5 s and a blank screen for 6 s (Fig. S1E). Subjects were instructed to silently read and memorize the sentence within the first 5 s and recite it in the next 5 s when the mouth cue appeared, followed by a 6 s rest with their eyes open.

In Scans #7 to #12, stimuli were presented auditorily through a pair of active noise canceling headphones. Subjects were instructed to listen to the speech attentively and keep their eyes open for the whole time. The speech of each sentence was synthesized using Google Cloud Text-to-Speech tool under default setting. The speech rate ranged from 2.9 to 3.1 syllables per second. For Scans #7 and #10, a speech with a visual cue (ear) was presented for 5 s, followed by a 11 s blank screen within each 16-s period (Fig. S1B). For Scans #8 and #11, a speech and a visual cue (ear and mouth) were presented for 5 s, followed by a 11 s blank period (Fig. S1D). Subjects were instructed to listen to the speech and repeat it right after it was heard. For Scans #9 and #12, a speech and a visual cue (ear) were presented for 5 s, followed by a visual cue (mouth) between 5-10 s and a 6 s blank screen between 10-16 s (Fig. S1F). Subjects were instructed to listen and memorize the speech during the first 5 s, and repeat the speech between 5-10 s.

Similar to the monolingual sessions, stimuli for the translation session were either visually or auditorily presented. Subjects would read silently or listen to the sentence in one language and orally translate into the other language when a mouth cue appeared. The translation session consisted of twelve 256-s functional scans, two scans for each of the six different tasks:

Scan #1: Reading the visually presented Chinese sentence silently, memorizing it, then orally translating it into English (L1-to-L2);

Scan #2: Listening to the auditorily presented Chinese sentence, memorizing it, then orally translating it into English (L1-to-L2);

Scan #3: Listening to five-digit number in Chinese, memorizing it, then orally translating it into English (L1-to-L2);

Scan #4: Reading the visually presented Chinese sentence silently, memorizing, then orally translating it into English (L1-to-L2);

Scan #5: Listening to the auditorily presented Chinese sentence, memorizing it, then orally translating it into English (L1-to-L2);

Scan #6: Listening to five-digit number in Chinese, memorizing it, then orally translating it into English (L1-to-L2);

Scan #7: Reading the visually presented English sentence silently, memorizing it, then orally translating it into Chinese (L2-to-L1);

Scan #8: Listening to the auditorily presented English sentence, memorizing it, then orally translating it into Chinese (L2-to-L1);

Scan #9: Listening to the five-digit number in English, memorizing it, then orally translating it into Chinese (L2-to-L1);

Scan #10: Reading the visually presented English sentence silently, memorizing it, then orally translating it into Chinese (L2-to-L1);

Scan #11: Listening to the auditorily presented English sentence, memorizing it, then orally translating it into Chinese (L2-to-L1);

Scan #12: Listening to the five-digit number in English, memorizing it, then orally translating it into Chinese (L2-to-L1).

The presentation paradigm for the visually presented stimuli was the same as that for Scans #3 and #6 of the monolingual sessions (Fig. 1E). The presentation paradigm for the auditorily presented stimuli was the same as that for Scans # 9 and #12 of the monolingual sessions (Fig. 1F).

### Experimental setup

To prevent head motion during tasks involving speaking, we made “The Phantom of the Opera” style masks molded to individual faces using thermoplastic sheets (1.6 mm H-board, Sun Medical Products Co., Ltd.). Each subject participated in a brief training session in an MRI simulator (Shenzhen Sinorad Medical Electronics Co., Ltd.) equipped with a motion sensor (MoTrak, Psychology Software Tools, Inc.). The subject would wear a mask and put on a pair of headphones, and practice keeping the head still while speaking in the MRI simulator. The head motion was monitored in real time by a sensor attached to the subject’s forehead. Auditory feedback (‘ding’ sounds) would be heard via the headphones whenever the head motion exceeded a threshold (1 mm in translation or 1° in rotation).

Before each fMRI session, the subject would wear a mask and earplugs, and put on a pair of MR-compatible noise-canceling headphones (OptoActive II, OptoAcoustics Ltd.). The subject would then lie supine with the head and headphones surrounded by deformable resin clay and silicone gel inside a head coil (Fig. S2A). An MR-compatible microphone (OptoActive II, OptoAcoustics Ltd.) was placed above the subject’s mouth (Fig. S2B, D), which allowed the subject to hear his or her own voice in real time via the headphones. A rear-mirror was mounted on top of the head coil, allowing the subject to view visual stimuli on an MR-compatible 40” LCD monitor (InroomViewingDevice, NordicNeuroLab AS) located 60 cm from the scanner bore (Fig. S2C, E). The total distance between the eyes and LCD monitor (via a mirror) was about 180 cm. The visually presented sentences subtended a maximum width of 39 cm on screen, resulting in a 12.4° horizontal field of view (6.2° eccentricity). An MR-compatible eye tracker (Eyelink 1000 Plus, SR Research Ltd.) was mounted behind the scanner bore, capturing the subject’s eye movements via the mirror (Fig. S2E, F). Both the mirror and the eye tracker were calibrated immediately after the subject was moved into the scanner bore. Visual and auditory stimuli were presented using Experiment Builder (SR Research Ltd.), which awaited a trigger signal (“s”) from SyncBox (NordicNeuroLab AS) before starting each functional scan.

### Behavioral recording

To ensure task compliance, speech output and eye movements were recorded in sync with functional scans. The subject’s voice was recorded into four soundtracks using the OptiMRI software (OptoAcoustics Ltd.) and analyzed using the Audacity ® software (https://audacityteam.org; Fig. S3). The group averages of subjects’ speech onset, offset, and duration are summarized in Table S3. For tasks that involved reading, a heat map was generated using EyeLink DataViewer (SR Research Ltd.) for the interest period (0 to 5 s within each 16 s trial) with fixation duration and fixation count (Fig. S4).

### Image acquisition

Functional and structural brain images were acquired using Siemens MAGNETOM Prisma 3T MRI scanner with a 32-channel head coil at the Centre for Cognitive and Brain Sciences, University of Macau. Each fMRI session consisted of twelve functional scans and 1 to 2 structural scans, which in total lasted for about 1.5 hours. Each functional scan was acquired using a blipped-CAIPIRINHA simultaneous multi-slice (SMS), single-shot echo planar imaging (EPI) sequence with the following parameters: acceleration factor slice = 5; interleaved ascending slices; TR = 1000 ms; TE = 30 ms; flip angle = 60°; 55 axial slices; field of view = 192×192 mm; matrix size = 64×64; voxel size = 3×3×3 mm; bandwidth = 2368 Hz/Px; 256 TR per image; dummy = 6 TR; effective scan time = 256 s. After the sixth functional scan, a set of T1-weighted structural images (alignment scan) was acquired using an MPRAGE sequence with the same center of slice groups and orientation as the functional images and with the following parameters: TR = 2300 ms; TE = 2.26 ms; TI= 900 ms; Flip angle = 8°; 256 axial slices; field of view = 256 x 256 mm; matrix size = 256×256; voxel size = 1×1×1 mm; bandwidth = 200 Hz/Px; scan time = 234 s. For the first session of each subject, a second set of structural images was acquired with parameters identical to the first set after the last functional scan.

### Data preprocessing

For each fMRI session, raw functional images were converted to the Analysis of Functional NeuroImages (AFNI) BRIK format. All BRIK files were registered with the first volume (target) of the seventh BRIK and corrected for motion using AFNI’s *3dvolreg* tool. The volume registration process yielded the time series of six degrees of freedom (three translational and three rotational motion parameters; Fig. S5). With head restraining measures, including custom-molded masks and deformable filling inside the head coil, no subject has shown major motion artifacts in functional images.

Bilateral cortical surfaces of each subject were reconstructed from the average of two sets of structural images using FreeSurfer 7.2 (*50*) (https://surfer.nmr.mgh.harvard.edu/). All motion-corrected functional images were registered with the alignment scan (structural images) in each fMRI session, and in turn registered with each subject’s cortical surfaces using FreeSurfer-compatible *csurf* (*33*) (https://pages.ucsd.edu/~msereno/csurf/ or https://mri.sdsu.edu/sereno/csurf/), which includes tools for functional image analyses as detailed below.

### Functional image analyses

For each BRIK file of a functional scan containing (*x*, *y*, *z*, *t*) = 64×64×55×256 data points, a 256-point discrete Fourier transform was applied to the time series *x_m_*(*t*) of each voxel *m* at location (*x*, *y*, *z*) by:

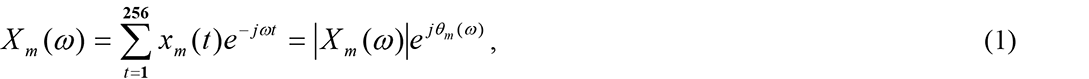

where *X*(*ω*) is the Fourier component at each frequency *ω* between 0-127 cycles per scan (cps), and |*X_m_*(*ω*)| and *θ_m_*(*ω*) represent the amplitude and phase angle respectively. The task frequency is defined as *ω_s_* (16 cps), and the remaining frequencies are defined as *ω_n_*. The signal and noise are defined as the Fourier components *X_m_*(*ω*) at frequencies *ω_s_* and *ω_n_* respectively. The statistical significance of signal-to-noise ratio in each voxel *m* is evaluated by an *F*-ratio (*29–32, 34*):

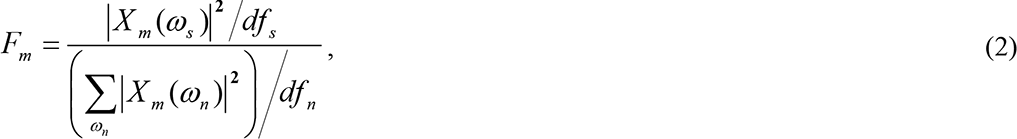

where *df_s_* = 2 and *df_n_* = 230 are the degrees of freedom of the signal and noise, respectively. The *P*-value in each voxel *m* is estimated by the cumulative distribution function *F*_(2, 230)_. A complex *F-value*, (*F_m_r_*, *F_m_i_*), incorporating both the *F*-statistic value and the phase angle, *θ_m_*(*ω_s_*), of each voxel was computed by *F_m_r_* = *f_m_* cos(*θ_m_*(*ω_s_*)) and *F_m_i_*= *f_m_* sin(*θ_m_*(*ω_s_*)), where *f_m_*is the square root of *F_m_*. Voxels containing strong periodic signals at the task frequency (*ω_s_* = 16 cps) with *F*_(2,230)_ > 4.7 (*P* < 0.01, uncorrected), *F*_(2,230)_ > 7.1 (*P* < 0.001, uncorrected), or *F*_(2,230)_ > 9.6 (*P* < 0.0001, uncorrected) were retained and their phase angles were color-coded in a range between 0−2π (0−16 s) and painted on each individual subject’s cortical surfaces for each scan using *csurf*. The complex *F-value*s of corresponding voxels *m* were vector-averaged (voxel-wise) across two scans *k* ={*1*, *2*} of the same task in each session for each subject *S* using:

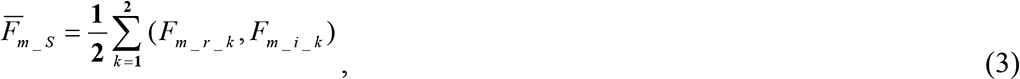

which was performed by the “Combine 3D Phase Stats” function of *csurf*. The resulting average complex *F*-values 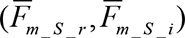 were then painted on individual subject’s cortical surfaces.

To find consistent spatiotemporal activation patterns across subjects, surface-based group averaging methods (*31–34, 51–53*) (*csurf* “Cross Sess SphereAvg” function) were used to resample all surface maps containing single-subject average complex *F*-values to a common spherical coordinate system for each hemisphere. The complex *F*-values at each vertex *v* on the common spherical surface were vector-averaged (vertex-wise) across all subjects (N = 21) using:

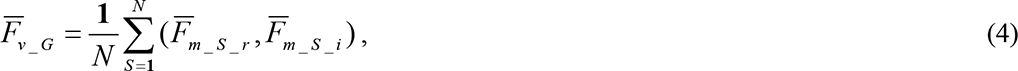

which yielded a map of group-average complex values 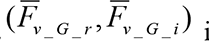 for each language task. Lastly, the resulting group-average complex *F*-values in the common spherical surface were projected back to the cortical surface of each hemisphere of a representative subject, which were displayed as single-subject phase-encoded activation maps using *csurf tksurfer* user interface (Figs. 1, 2).

### Static and dynamic brain ‘weather’ maps

To unfold the complex spatiotemporal patterns in phase-encoded maps, we created isophase contour maps to illustrate the directions and wavelengths of traveling waves on flattened cortical surfaces (Fig. 3). Vertices containing the same range of phase angles (i.e., isophase) were painted with white ‘surf line’ contours by setting the $phasecontourflag = 1 in *csurf tksurfer* (Fig. 3A to D). To find the propagation paths of traveling waves, a vector field map of phase gradient (cf. maps of wind or ocean current patterns) was estimated using a nearest neighbor vertex basis on the flattened cortical surface (*tksurfer compute_surfgrad* tool). The resulting principal phase gradient directions are indicated by arrows (paths) in the leftmost panel of each movie frame sequence in Fig. 3 (E to P).

In addition to static maps of isophase contours and phase gradient, we created a ‘brainstorm’ movie for each language task using the *tksurfer phasemovie.tcl* script to illustrate the dynamic traveling waves (cf. animated ‘rainbands’ of an impending storm; see Movies S1 to S18). Each frame of a ‘brainstorm’ movie shows surface vertices containing periodic activations with phase angles falling in a moving window (range = 9°; step = 4.5°, equivalent to 0.2 s/frame) at each moment. The isophase activation patterns were rendered in phase-encoded colors using the colorbar in Figs. 1 and 2. In total, each ‘brainstorm’ movie contains 80 frames of spatiotemporal traveling waves within a period of 0−16 s ([0, 2π] or [0°, 360°]).

Multilayer brain ‘weather’ maps were constructed to facilitate interpretation of the complex spatiotemporal patterns of traveling waves and comparison across different language tasks (Fig. 4). The bottom layer (Fig. 4A) contains flattened cortical surfaces of the left and right hemispheres reconstructed from the structural brain images of a representative subject. This layer shows the anatomy (cf. terrain) of the cortical surface, with dark-gray regions representing sulci (cf. valleys) and light-gray regions representing gyri (cf. ridges). To navigate in ‘uncharted territory’ on the cortical surface, a recent brain atlas outlining the latest parcellation of topological visual, auditory, and somatomotor maps(*33*) (https://pages.ucsd.edu/~msereno/csurf/fsaverage-labels/) was mapped onto the cortical surfaces of this representative subject using the *csurf tksurfer annot2roi.tcl* script (Fig. 4A). The contours (gray) of topological areas were overlaid on flattened cortical surfaces in a way similar to the ‘county’ lines on a weather forecast map. This second layer was then used as a reference map (e.g., ‘landmarks’ of regions of interest) to guide the interpretation of functional activations on the maps of unimodal, multimodal, crossmodal, and cross-language tasks. To this end, we overlaid six layers of language maps containing contours taken from Fig. 1, which show the extent of brain activations (cf. clouds in a satellite image) in L1 or L2 reading-only (unimodal), listening-only (unimodal), and speaking (multimodal) tasks (Fig. 4B). We then illustrated the paths of surface traveling waves (cf. storm paths) during unimodal reading or listening tasks (black arrows in Fig. 4B) and during a crossmodal L1 reading-memorizing-reciting task (yellow arrows in Fig. 4C, redrawn from Fig. 3C).

As traveling waves ‘escaped’ from the topological maps, they made ‘landfall’ in language-specific and domain-general cognitive regions, which caused transient ‘storm surges’ in these regions. We selected nine sROIs, including rMFG, dlPFC, IPL, dmFAF, AIC, Spt, ATL, STS, and ITG (magenta contours in Fig. 4B, C), to compare the surge profiles of unimodal, multimodal, crossmodal, and cross-language tasks within each sROI on the group-average maps (Fig. 4D, Fig. S6). For each task, a surge profile was estimated from the distribution of group-average complex *F*-values (*F_v_G_r_*, *F_v_G_i_*) of all vertices within each sROI using circular statistics (*34, 54*). First, the phase angle of each vertex *v* was obtained by *θ_v_G_* = atan2 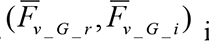 in Matlab. Second, the 16-s period was divided into 80 equally spaced bins in [0°, 360°] or [0, 2π]. A total of *V* vertices were found with phase angles *θ_v_G_* falling within the moving window centered at each bin *d* (range = 9°; step = 4.5°, equivalent to 0.2 s/bin). The complex *F*-values of *V* vertices in the moving window *d* were averaged using:

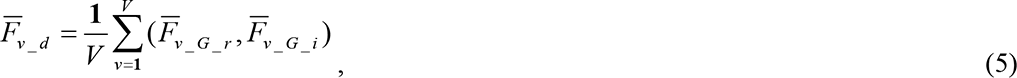

The magnitude of the average complex *F*-value was then obtained by:

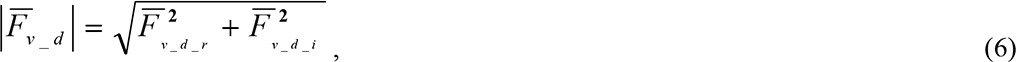

A *P*-value was estimated for each 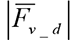 using the cumulative distribution function *F*_(2, 230)_. Lastly, the height of surge is computed by -log_10_(*P*-value) as shown in the y-axis of each subplot in Fig. 4D and Fig. S6. The resulting surge profiles reveal the arrival time, ending, latency, and peak magnitude of traveling waves in each sROI (Fig. 4C, D).

### University of Macau Brain Atlas (UMBA)

In this study, functional activation maps of unimodal, multimodal, crossmodal, and cross-language tasks were displayed on the same cortical surfaces, which enable direct comparison of spatiotemporal traveling wave patterns across languages and tasks on multilayer surface-based functional brain atlases (*32, 33, 55*). To share these results with other researchers, we constructed an open access website, the University of Macau Brain Atlas (UMBA), incorporating all maps and movies in the current study (Movie S19). This online atlas (available once this manuscript is accepted by a peer-reviewed journal) will undergo continuous updates with new layers of maps and movies of multiple populations, languages, and tasks from our ongoing studies involving more than 150 subjects.

The design of multilayer user interface in UMBA drew on concepts from Google Earth (Movie S20; https://earth.google.com/web/) and weather forecast websites (e.g., time-lapse view of moving rainbands on https://www.wunderground.com/wundermap/). The UMBA website was developed using HTML, CSS, and Javascript languages. The Vue.js frontend JavaScript framework (version 2.x; libraries: vuetify, vue-i18n) was used for ‘responsive reactive design’ for the user interface. Built on a pair of flattened cortical surfaces, each of the following layers can be made visible or hidden according to the user’s preference: (1) topological maps(*33*); (2) contours of topological areas; (3) annotation of major sulci; (4) three layers of maps of reading, listening, and speaking tasks for each language (6 layers in total); (5) contours of sROI; (6) paths of traveling wave; (7) labels (landmarks) of brain areas and sROI; (8) phase-encoded activation maps for each task and language; (9) ‘brainstorm’ (traveling wave) movie for each task and language. A demonstration of UMBA interface and multilayer maps is shown in Movie S19.

## Acknowledgments

We thank Yi Tang for participant recruitment and assistance with fMRI experiments; Chi Hang Choi and Aotong Li for the design and construction of UMBA website; Aotong Li and several other research assistants for data preprocessing and artwork.

## Funding

University of Macau Development Foundation grant EXT-UMDF-014-2021; University of Macau grants CPG2023-00016-FAH, CRG2021-00001-ICI, CRG2020-00001-ICI, SRG2019-00189-ICI; Macau Science and Technology Development Fund FDCT 0001/2019/ASE; U.S. National Institute of Health grant R01 MH081990 (MIS, RSH)

## Author Contributions

VLCL, TIL, MIS, DL and RSH contributed to the conceptualization and the methodology of the research as well as writing and reviewing the paper. TIL, CTL, LL, CUC and RSH collected and analyzed the data. VLCL, DL and RSH contributed to other aspects of the project including fund acquisition and supervision.

## Competing interests

Authors declare that they have no competing interests.

## Supplementary Figures

**Figure S1.**
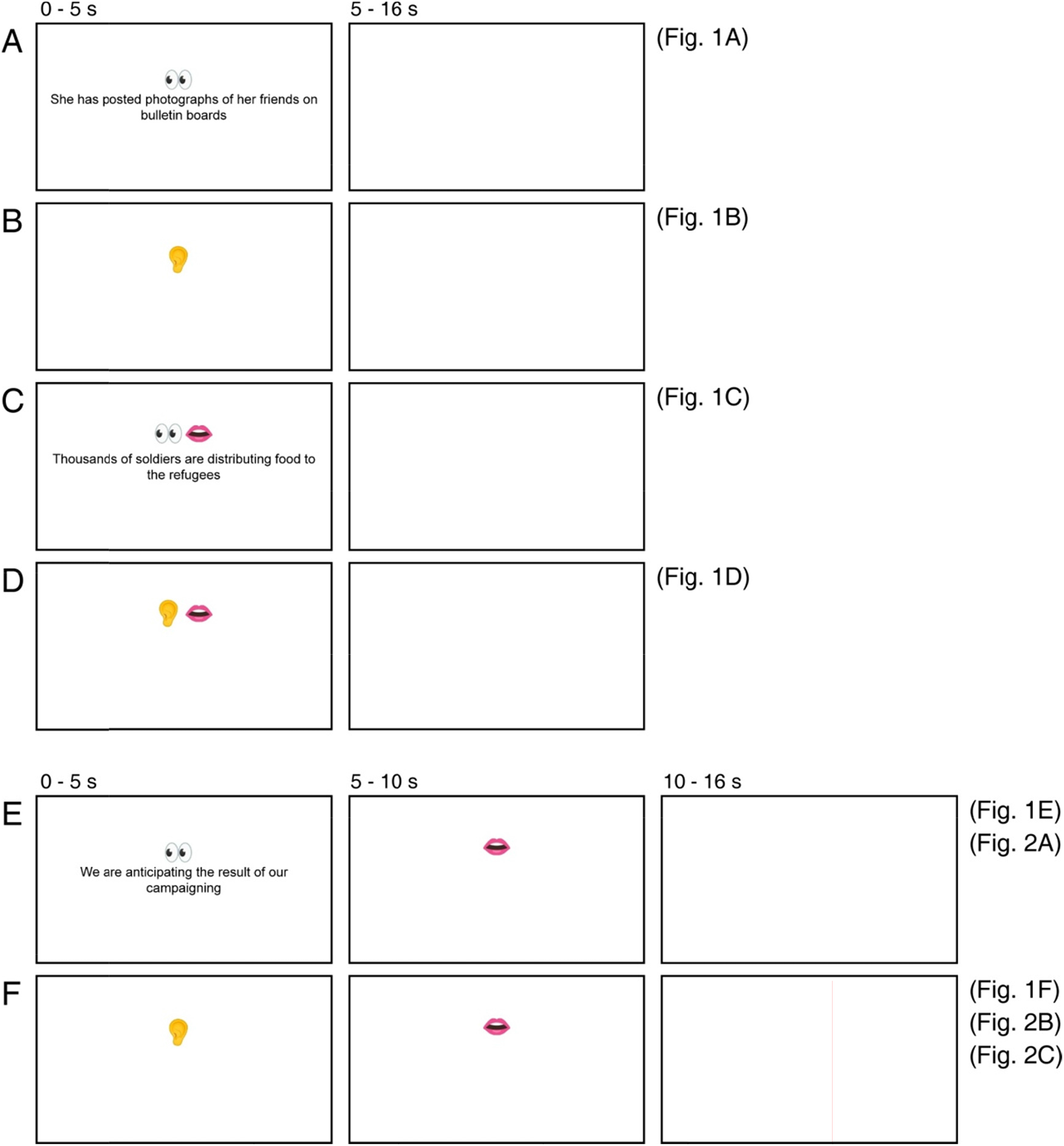
Experimental paradigms and stimuli. (**A**) Read**i**ng only tasks, corresponding to Fig. 1A. (**B**) Listening only tasks, corresponding to Fig. 1B. (**C**) Reading-aloud tasks, corresponding to Fig. 1C. (**D**) Shadowing tasks, corresponding to Fig. 1D. (**E**) Reading-me**m**orizing-reciting and reading-memorizing- translating tasks, corresponding to Figs. 1E and 2A. (**F**) For listening-memorizing-reciting and listening- memorizing-translating tasks, corresponding to Figs. 1F, 2B and 2C. In (A) to (D), visual or auditory stimuli were presented between 0-5 s, followed by a blank screen between 5-16 s. In (E) and (F), visual or auditory stimuli were screen between 5-16 s. resented between 0-5 s, followed by s**p**eech output between 5-10 s and a blank [Figure not shown in this preprint due to the inclusion of photographs of a human subject.]

**Figure S2.** Experimental setup. Figure not shown in this preprint due to the inclusion of photographs of a human subject. (**A**) Subject wearing a “The Phantom of the Opera” style mask and a pair of noise-canceling headphones, with resin clay filled around the head and headphones to reduce head motion and scanner noise. (**B**) A microphone in front of the subject’s mouth. (**C**) A rear-view mirror mounted above the 32-channel head coil. (**D**) Acoustic foams placed inside the bore for scanner noise reduction. (**E**) MR-compatible LCD monitor and eye tracker behind the scanner. (**F**) A close-up view of the eye tracker.

**Figure S3.**
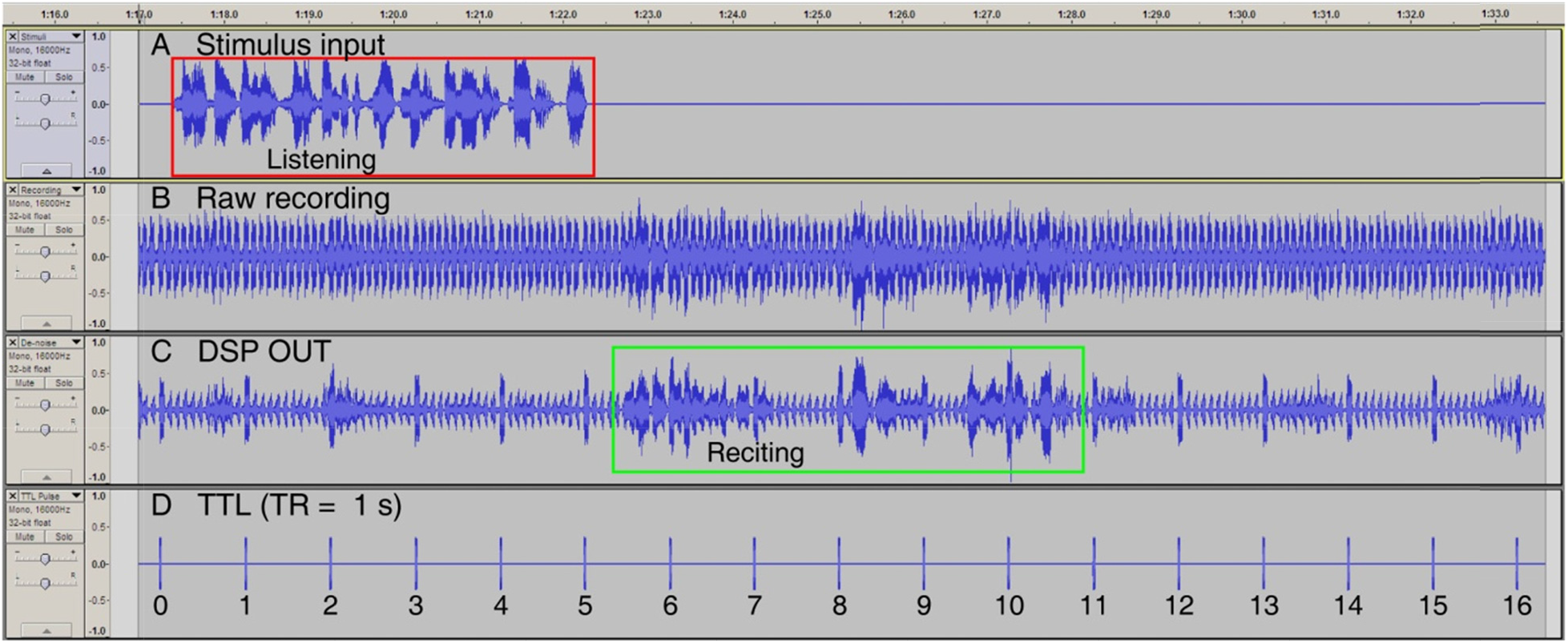
Audio recording. Sample soundtracks of a single trial in an L2 listening-memorizing-reciting task. (**A**) Recording of audio stimulus input. (**B**) Raw recordi**n**g: speech output recorded with unprocessed MRI noise. (**C**) Digital signal processing (DSP) OUT: audio **o**utput after applying real-time noise cancellation to the raw recording in (B). (**D**) Transistor–transistor logic (TTL) pulse: timestamps recorded at the beginning of each TR. The screenshot of recordings was created usi**n**g Audacity® (Audacity® software is copyright © 1999-2021 Audacity Team. The name Audacity® is a registered trademark. https://www.audacityteam.org/about/citations-screenshots-and-permissions/).

**Figure S4.**
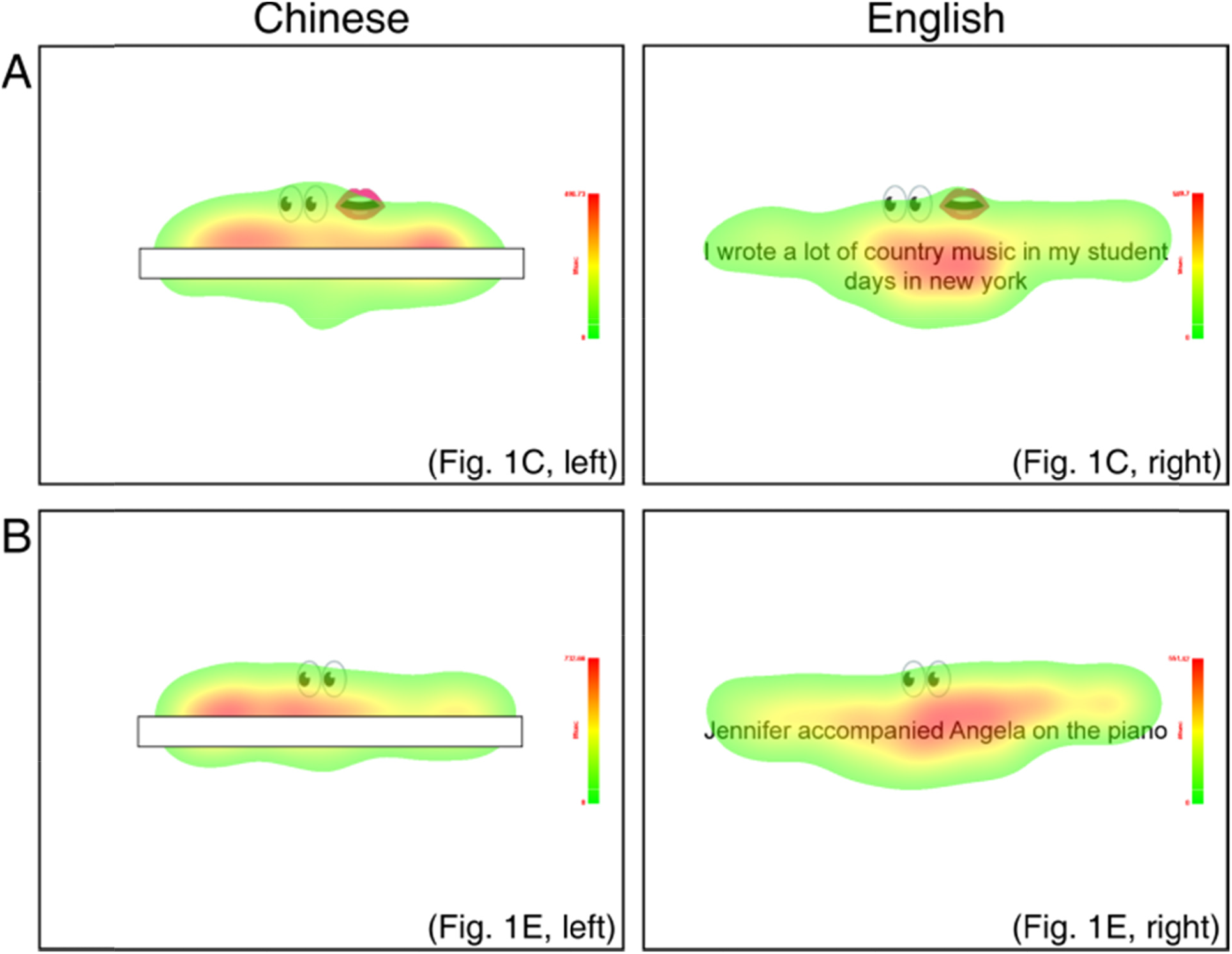
Eye-tracking heat maps. Results of eye-tracking for tasks involving reading in one representative subject. (**A**) Heat maps for reading-aloud tasks (see Fig. 1C). (**B**) Heat maps for reading- memorizing-reciting tasks (see Fig. 1E). (Left panels) Chinese, (Right panels) English. The English translation of two example Chinese sentences presented in the boxes are: (A) *The suspect has already left the crime scene*, and (B) *Successful enterprises invested on this premium product*.

**Figure S5.**
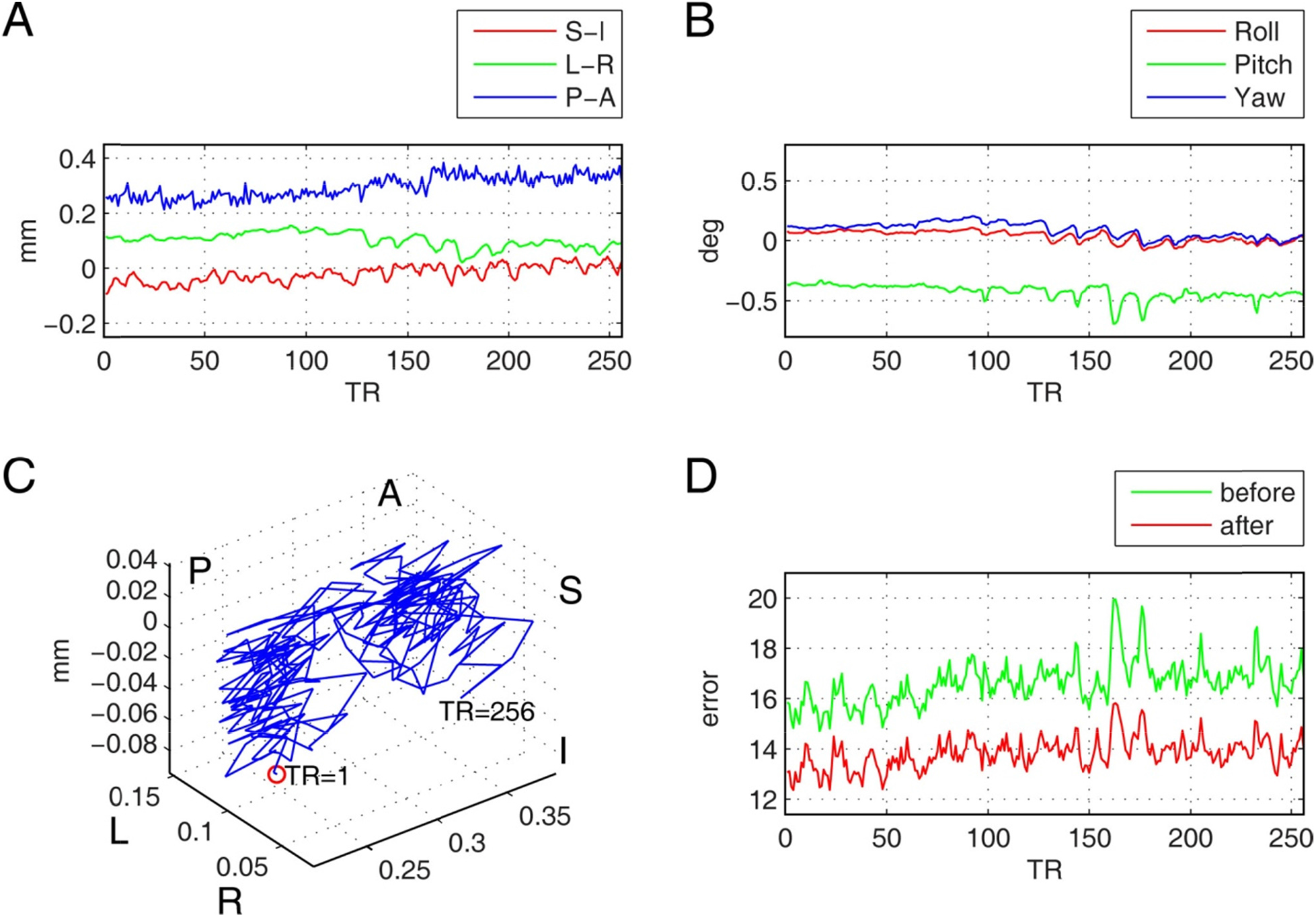
Head motion trajectory. Results of head motion correction for a representative scan that involved speaking. (**A**) Head translation (mm) in three axes: superior-inferior (S-I), left-right (L-R), and posterior-anterior (P-A) directions. (**B**) Head rotation (degree) in three directions. (**C**) 3D trajectory of head motion from TR=1 to TR=256. (**D**) Time series of root-mean-square difference (error) between each volume and the reference volume, before and after motion co**r**rection.

**Figure S6.**
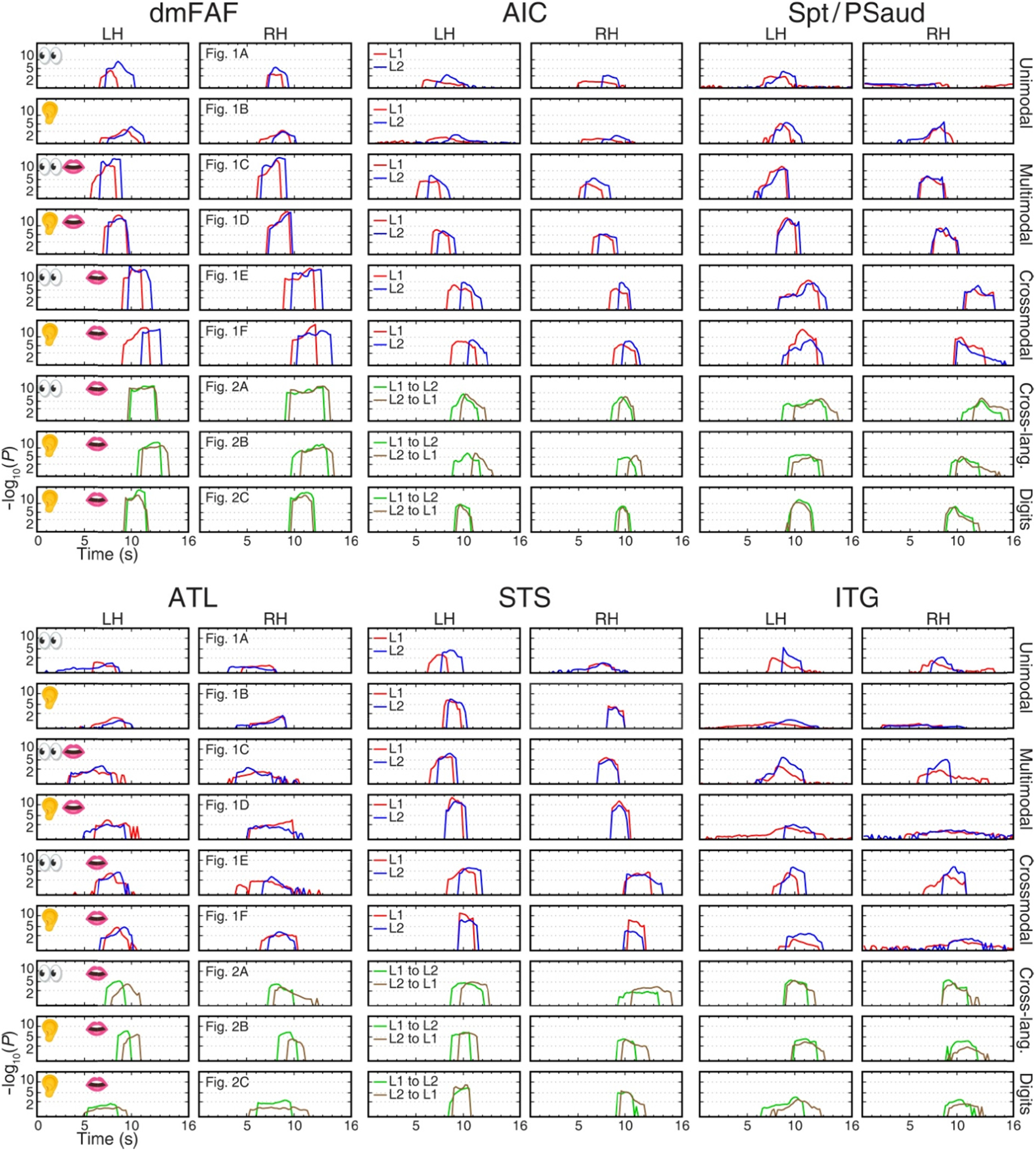
Surge pro iles of traveling waves in six sROIs. The figure legends follow Fig. 4D.

**Table S1.** **English translation of sixteen Chinese (L1) example sentences.**

|  |
| --- |
| Successful enterprises invested in this premium product. |
| Elementary school students visited this museum. |
| The United Nation successfully tackled the severe food crises. |
| The central government is distributing epidemic prevention guidelines. |
| Indonesian leaders have signed an agreement. |
| The botanists patiently explained the relevant issues. |
| My parents have increased my living expenses this semester. |
| Archaeologists stumbled upon prehistoric remains. |
| A massive lightning bolt struck the top of Oriental Pearl Tower. |
| The Bank of China has hired professional statisticians. |
| Graduate students were patiently collecting experimental data. |
| My dad gave me a beautifully wrapped gift. |
| This newly developed drug has reduced the mortality rate of the disease. |
| The zoo staff is feeding a lion cub. |
| The immigration officers carefully check all identification cards. |
| This naughty child caught many bugs. |

**Table S2.**
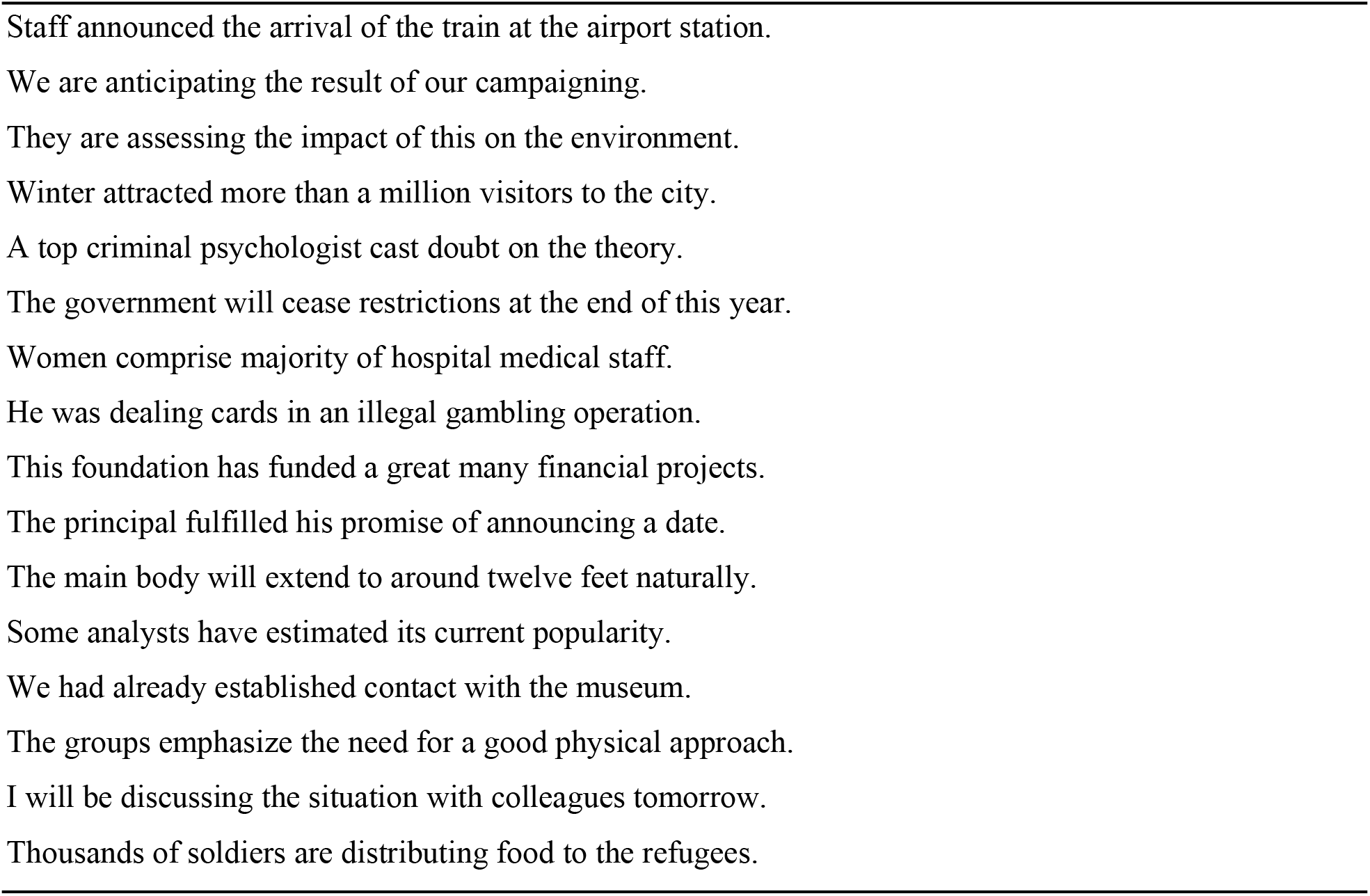
**Sixteen English (L2) example sentences.**

**Table S3.**
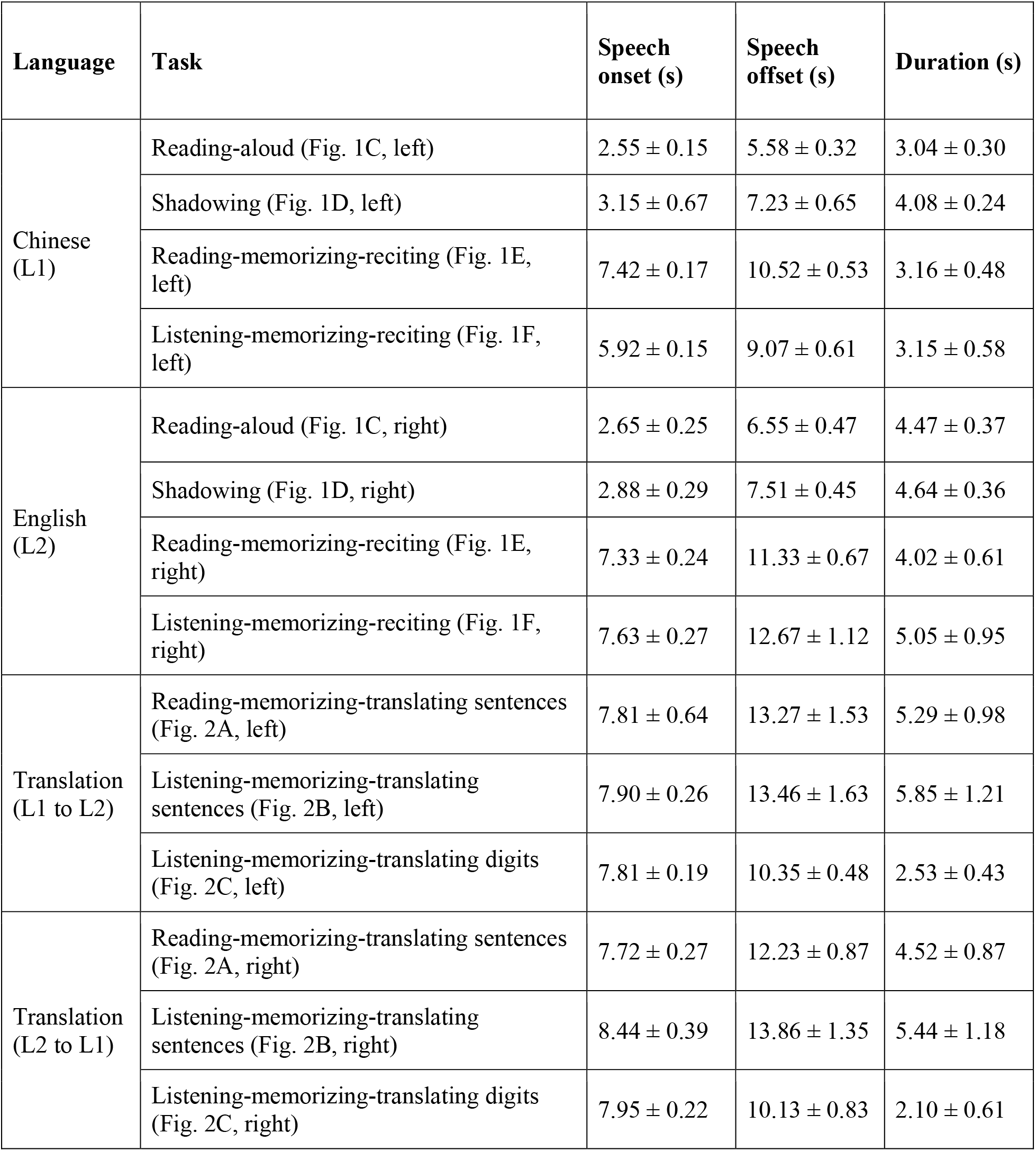
**Group average timing and standard deviation of recorded speech output.** Data from the same task were averaged across 21 subjects.

**Table S4.**
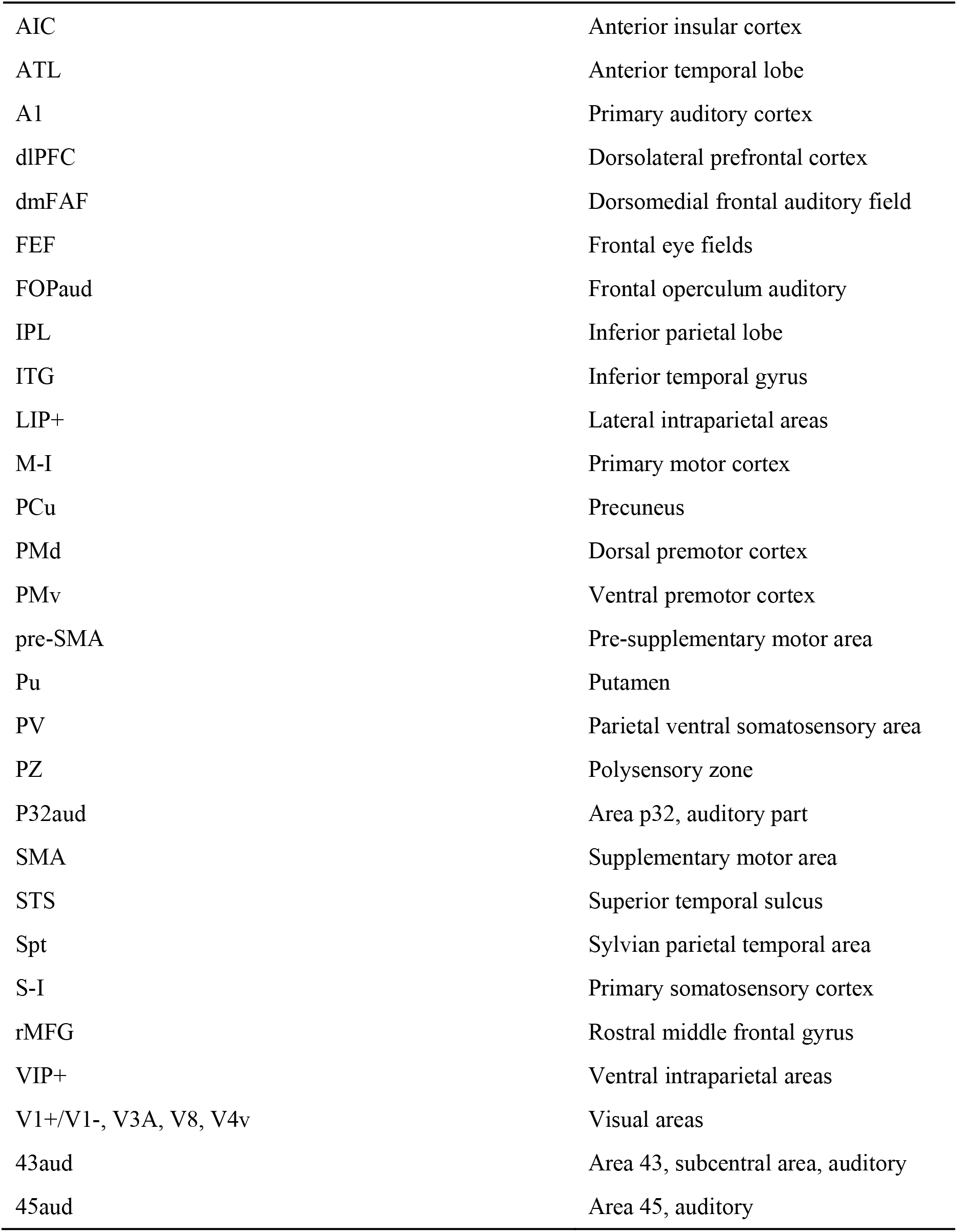
**Abbreviations of cortical areas in alphabetical order.**

## Supplementary Videos

**Movie S1.**
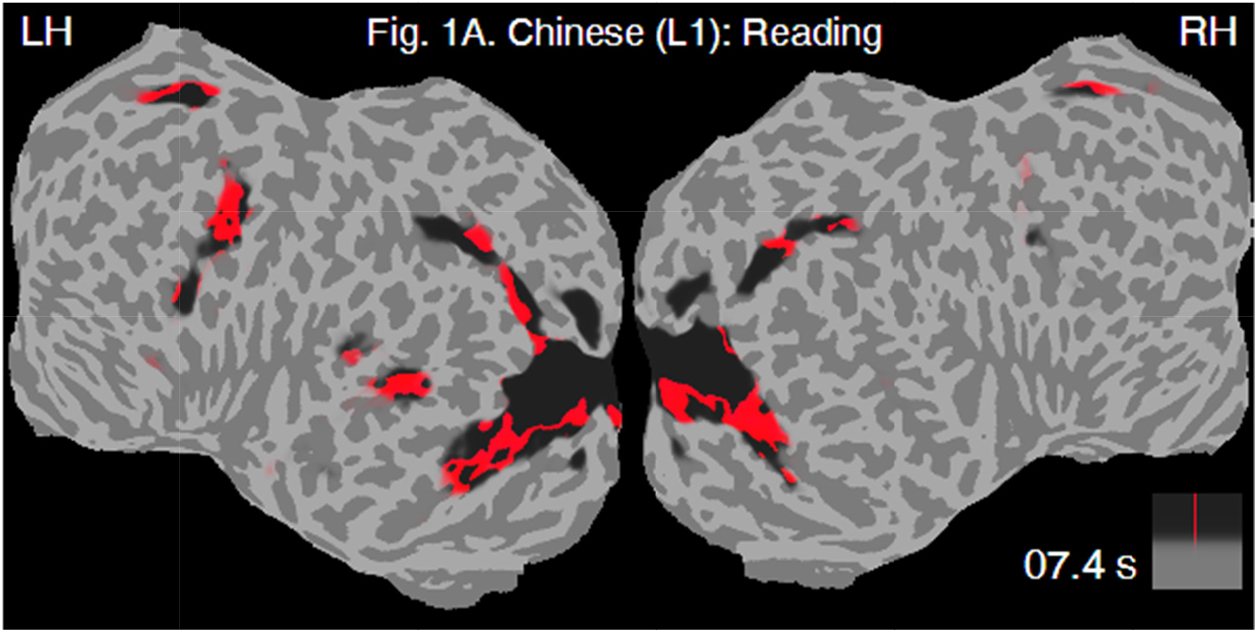
Brainstorms induced by a silent reading task in L1.

**Movie S2.**
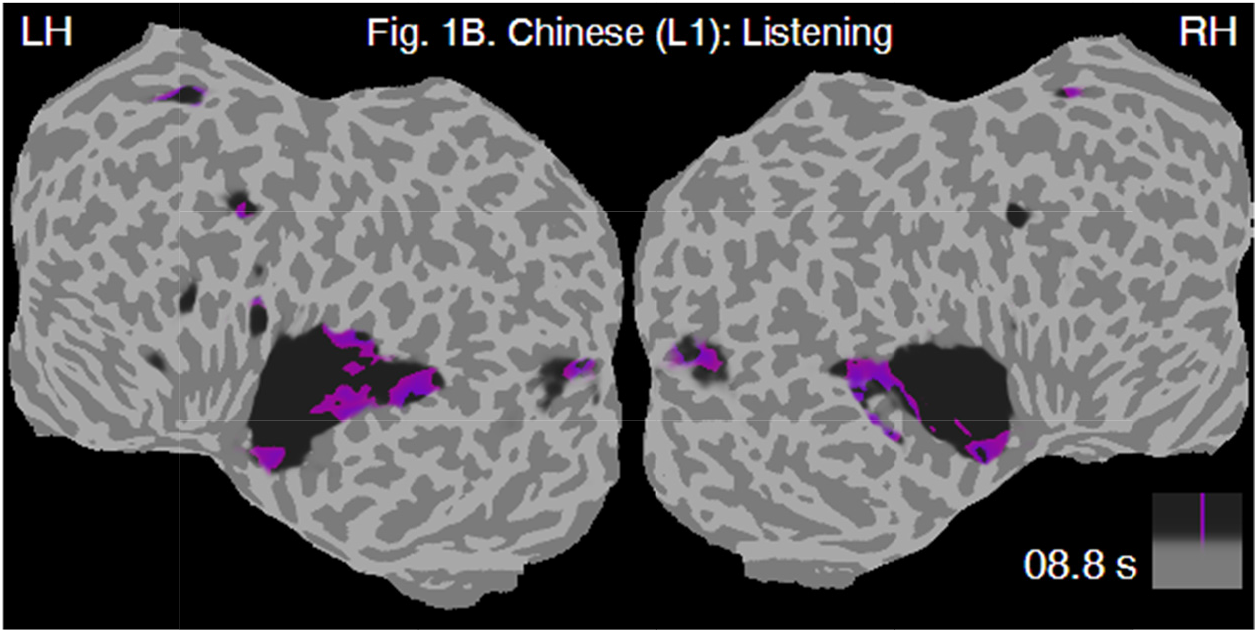
Brainstorms induced by a listening task in L1.

**Movie S3.**
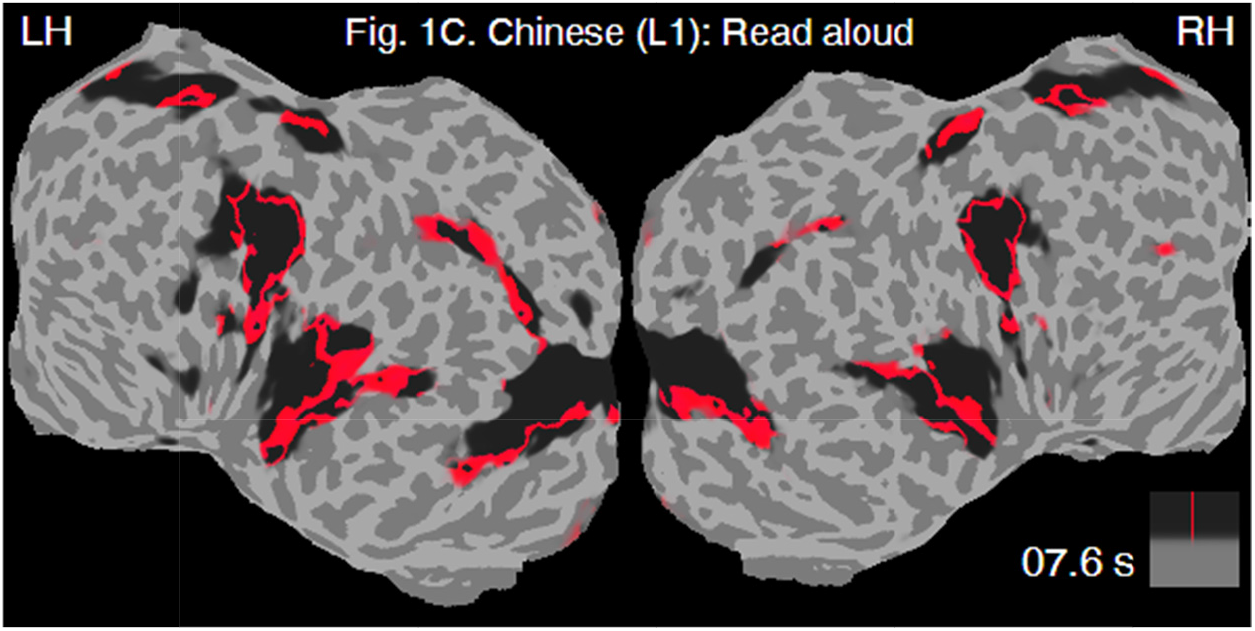
Brainstorms induced by a reading-aloud task in L1.

**Movie S4.**
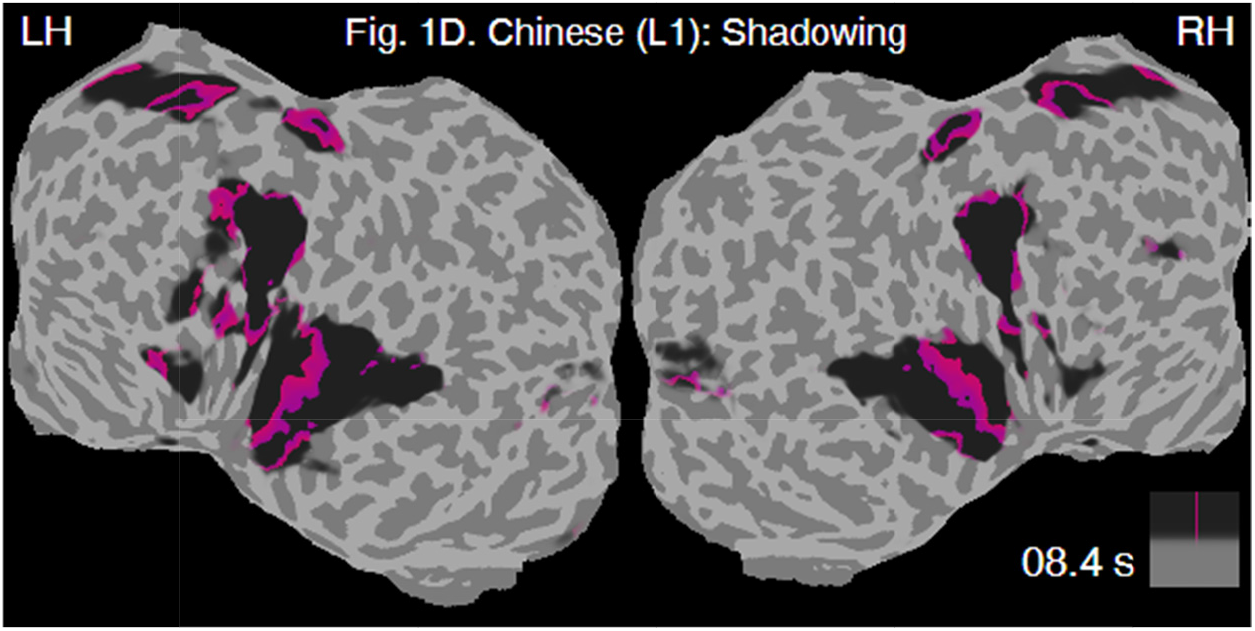
Brainstorms induced by a shadowing task in L1.

**Movie S5.**
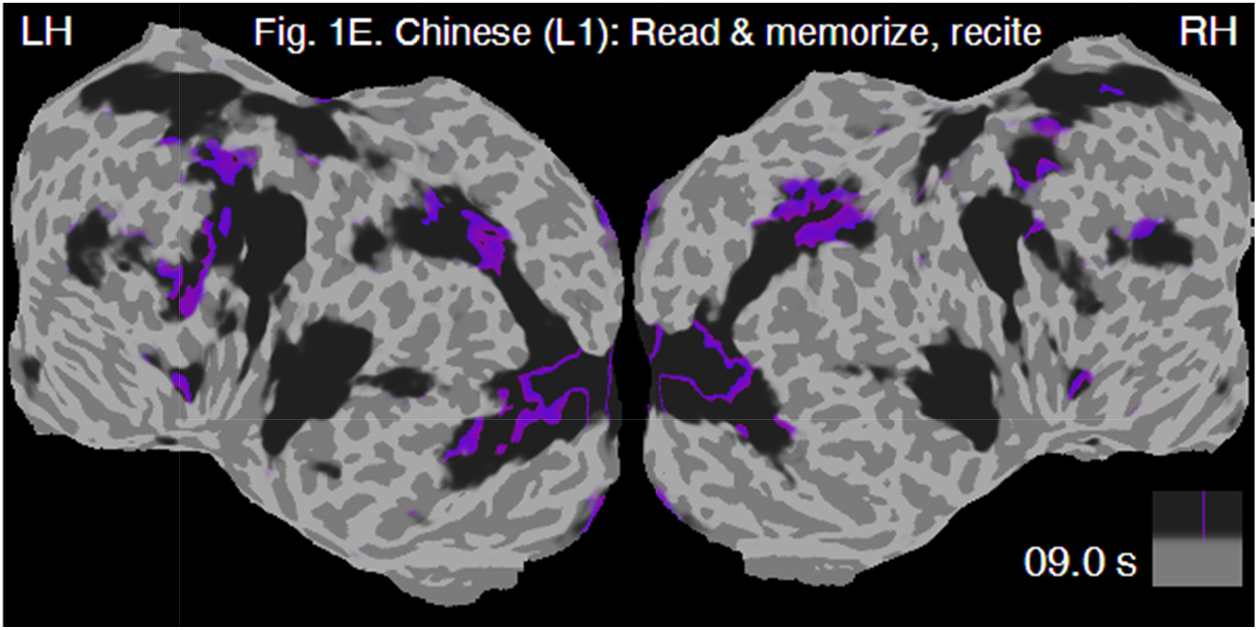
Brainstorms induced by a reading-memorizing-reciting task in L1.

**Movie S6.**
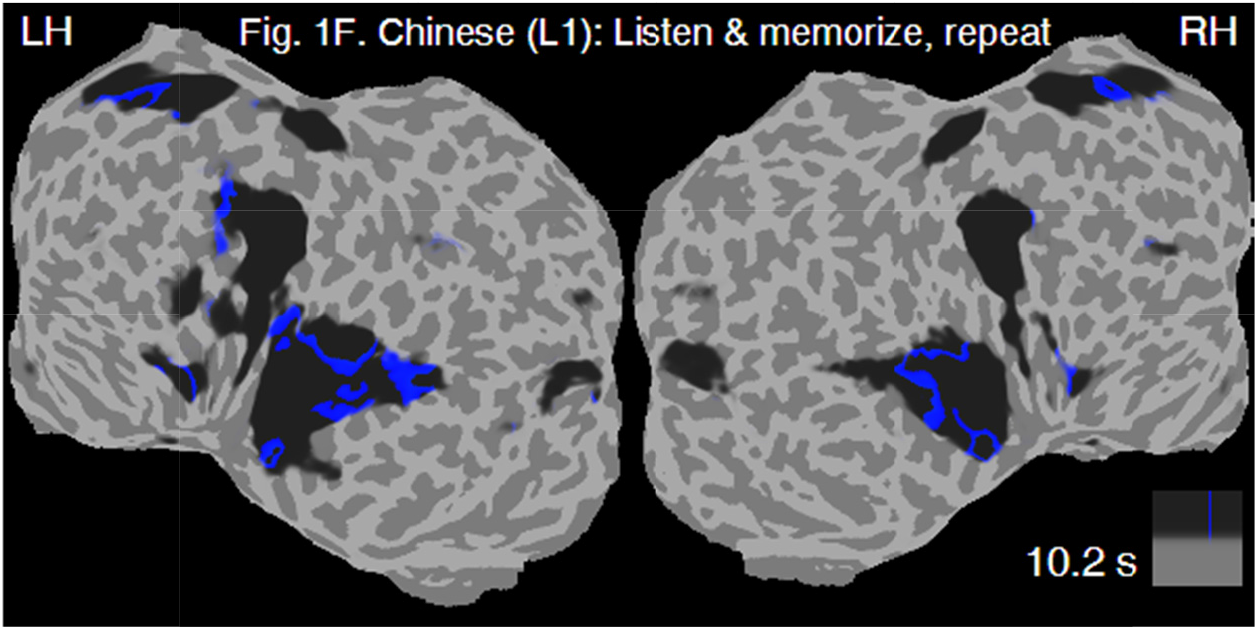
Brainstorms induced by a listening-memorizing-reciting task in L1.

**Movie S7.**
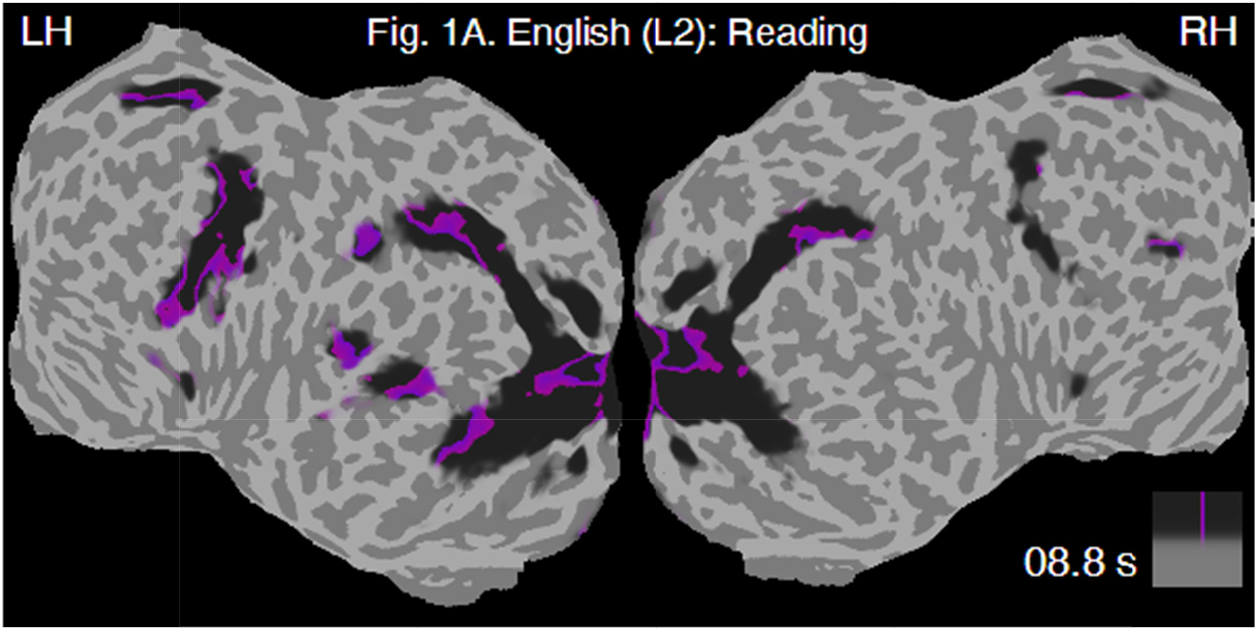
Brainstorms induced by a silent reading task in L2.

**Movie S8.**
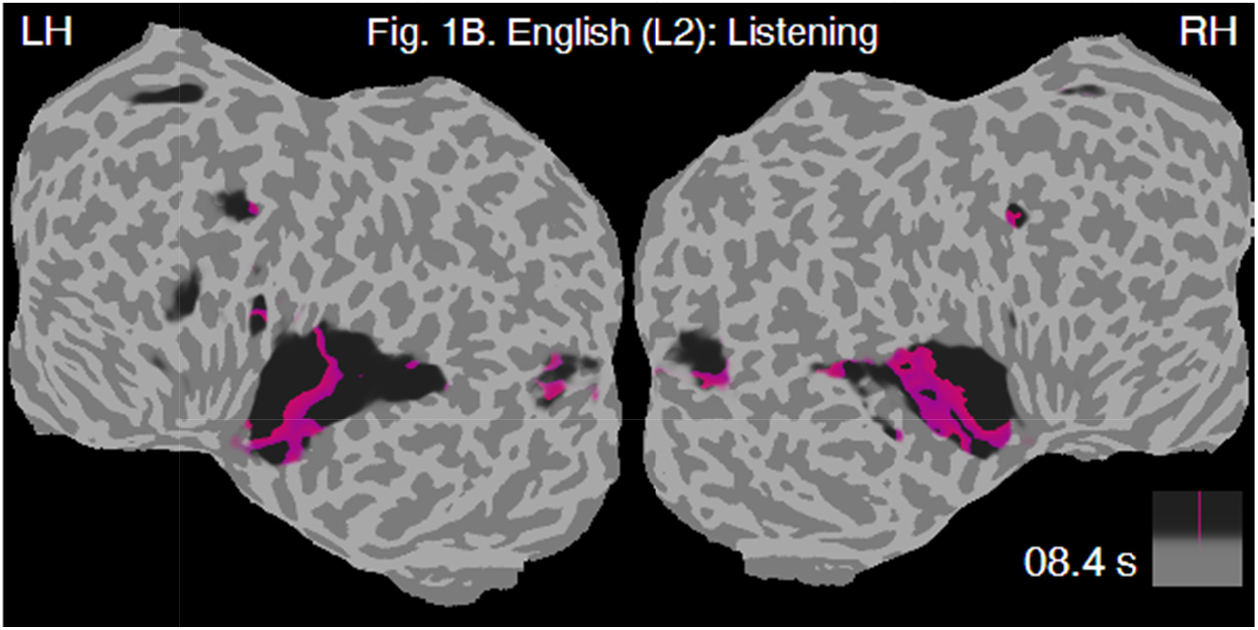
Brainstorms induced by a listening task in L2.

**Movie S9.**
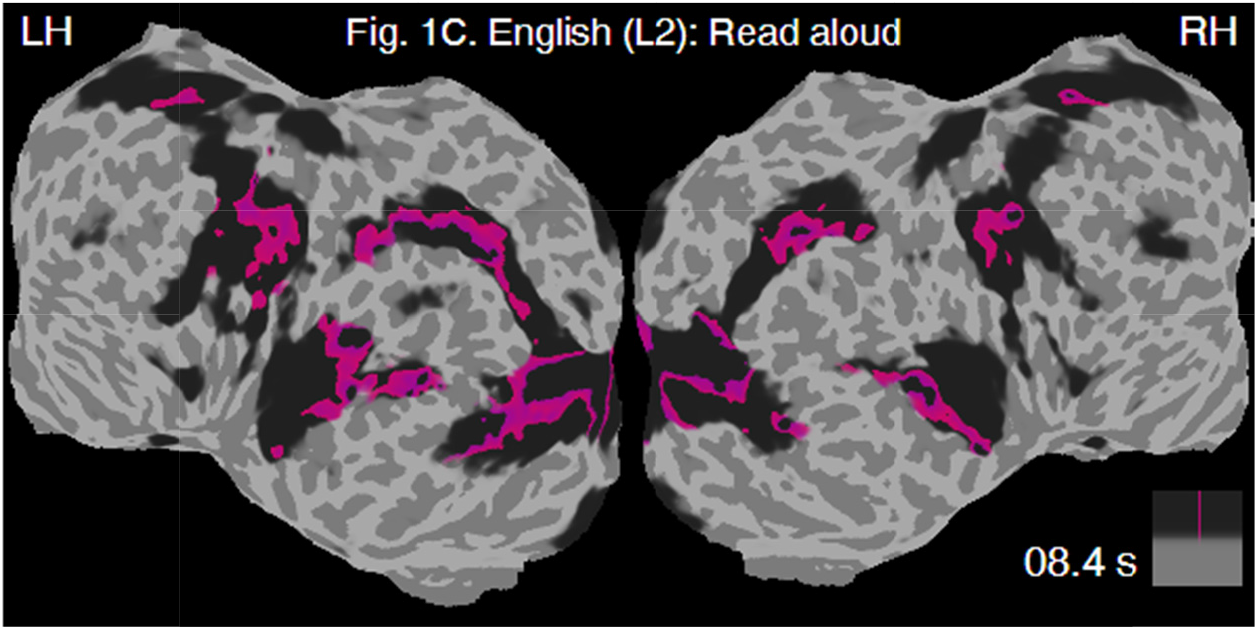
Brainstorms induced by a reading-aloud task in L2.

**Movie S10.**
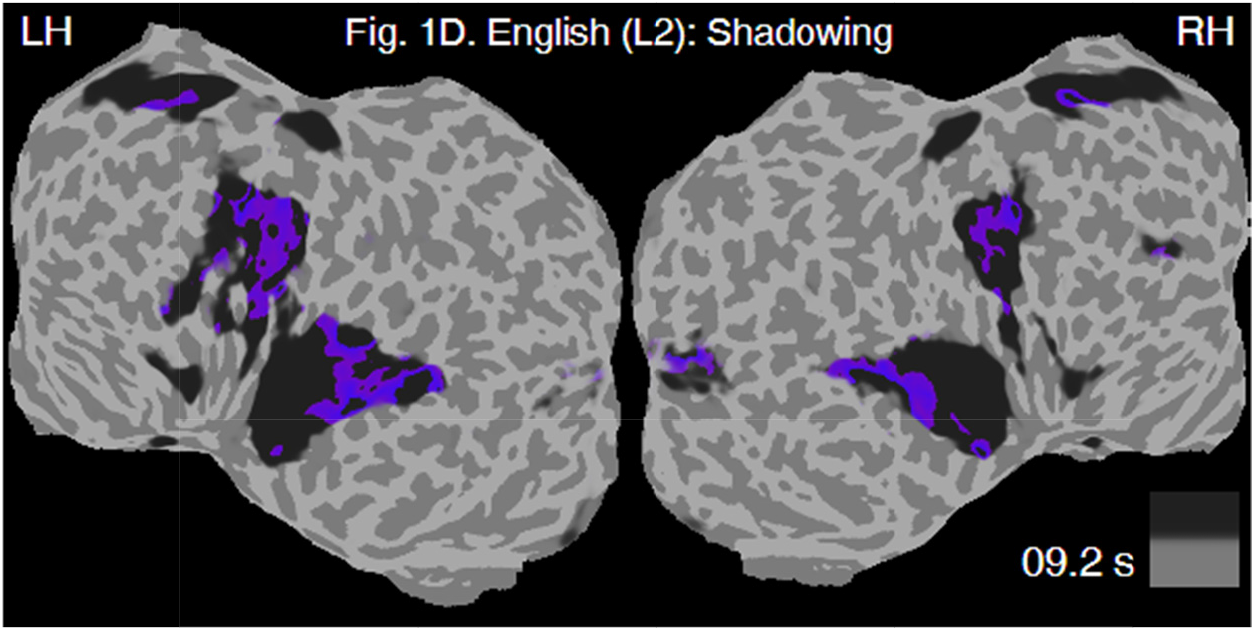
Brainstorms induced by a shadowing task in L2.

**Movie S11.**
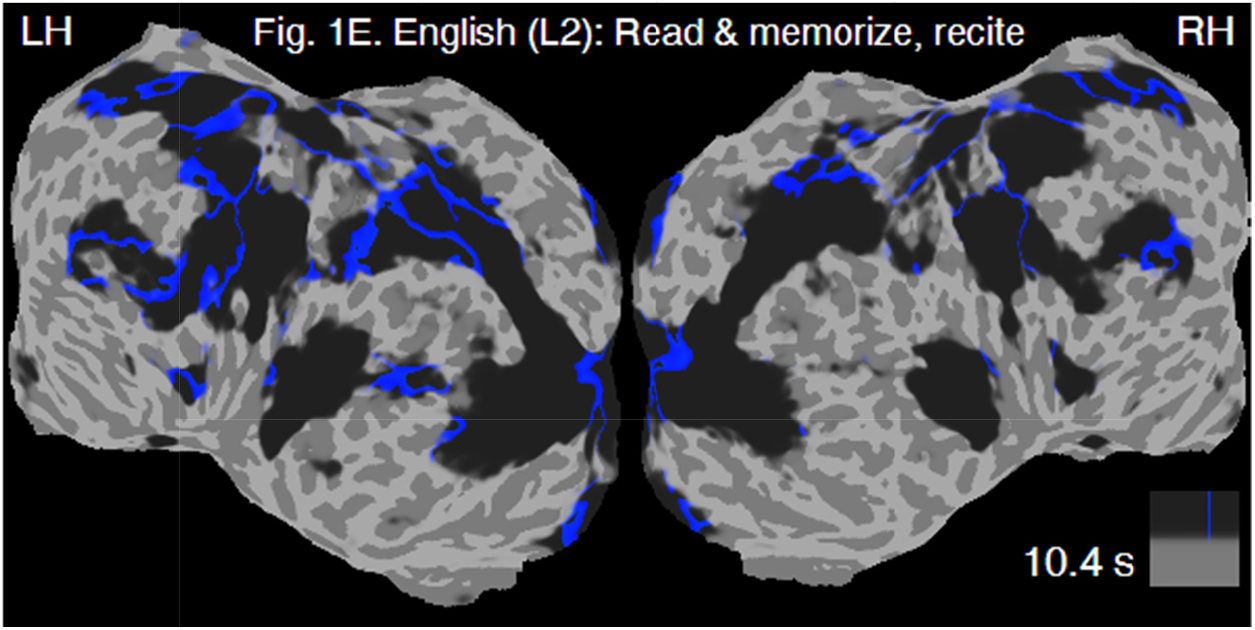
Brainstorms induced by a reading-memorizing-reciting task in L2.

**Movie S12.**
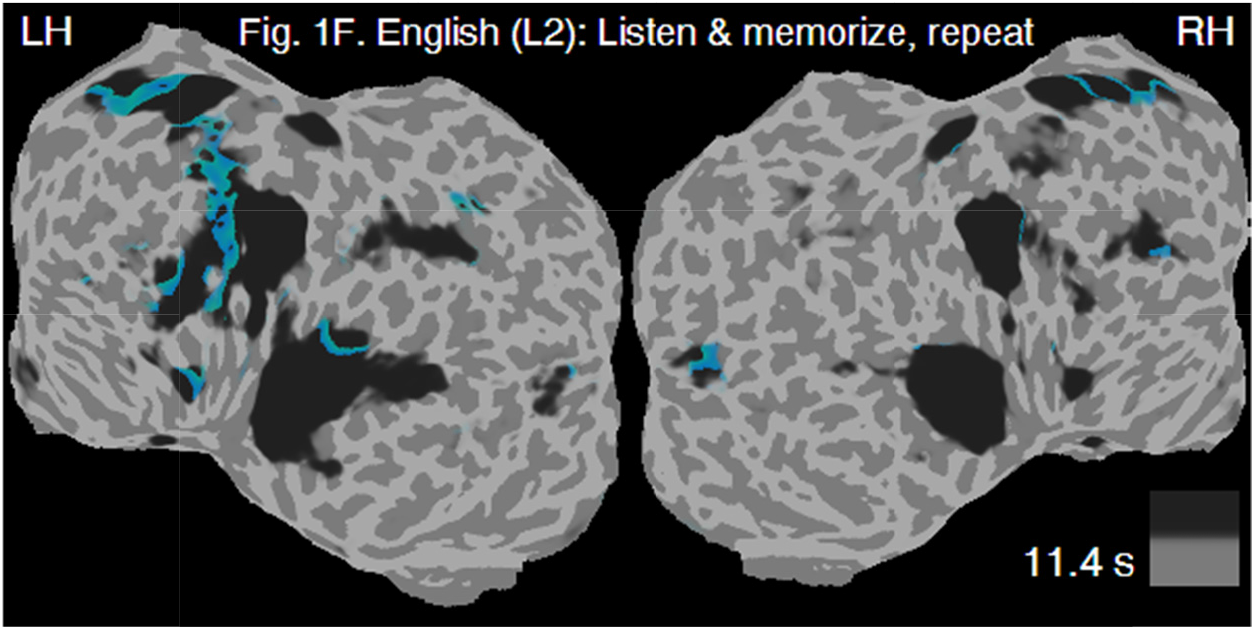
Brainstorms induced by a listening-memorizing-reciting task in L2.

**Movie S13.**
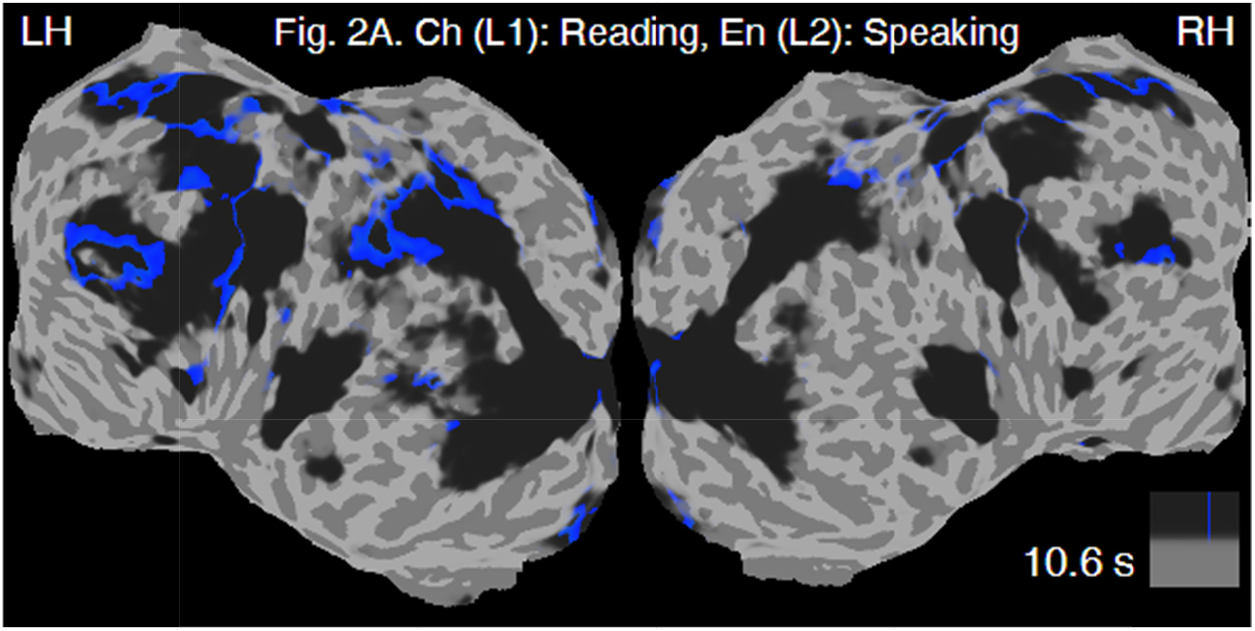
Brainstorms induced by an L1-to-L2 sight interpreting task.

**Movie S14.**
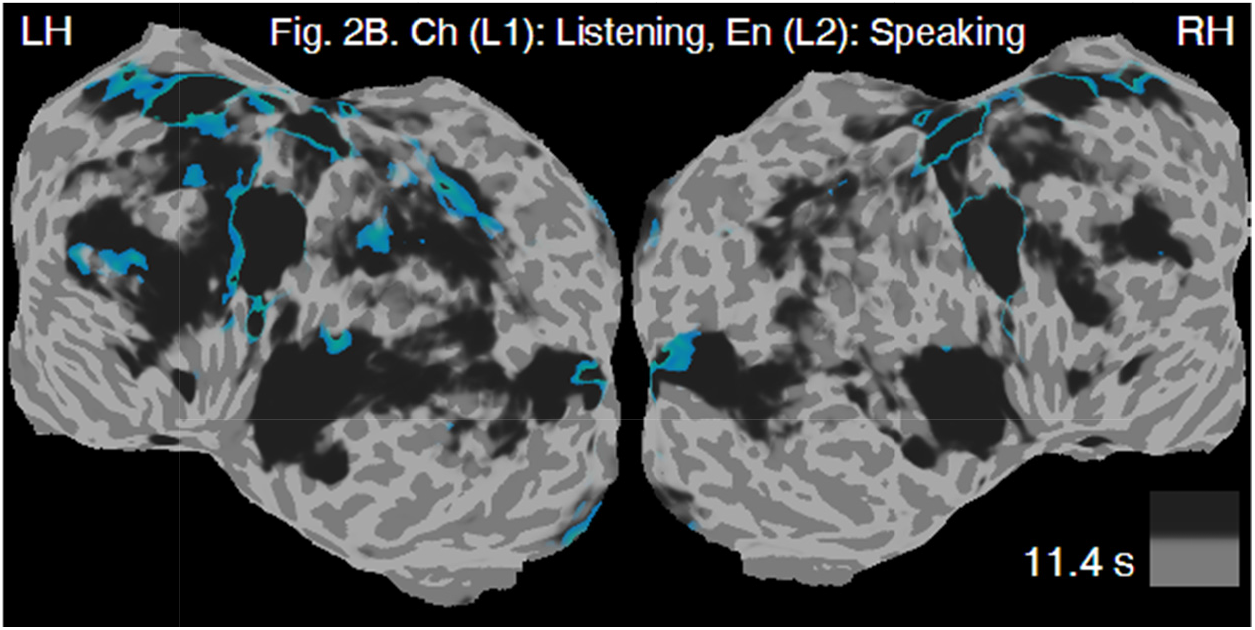
Brainstorms induced by an L1-to-L2 consecutive speech interpreting task.

**Movie S15.**
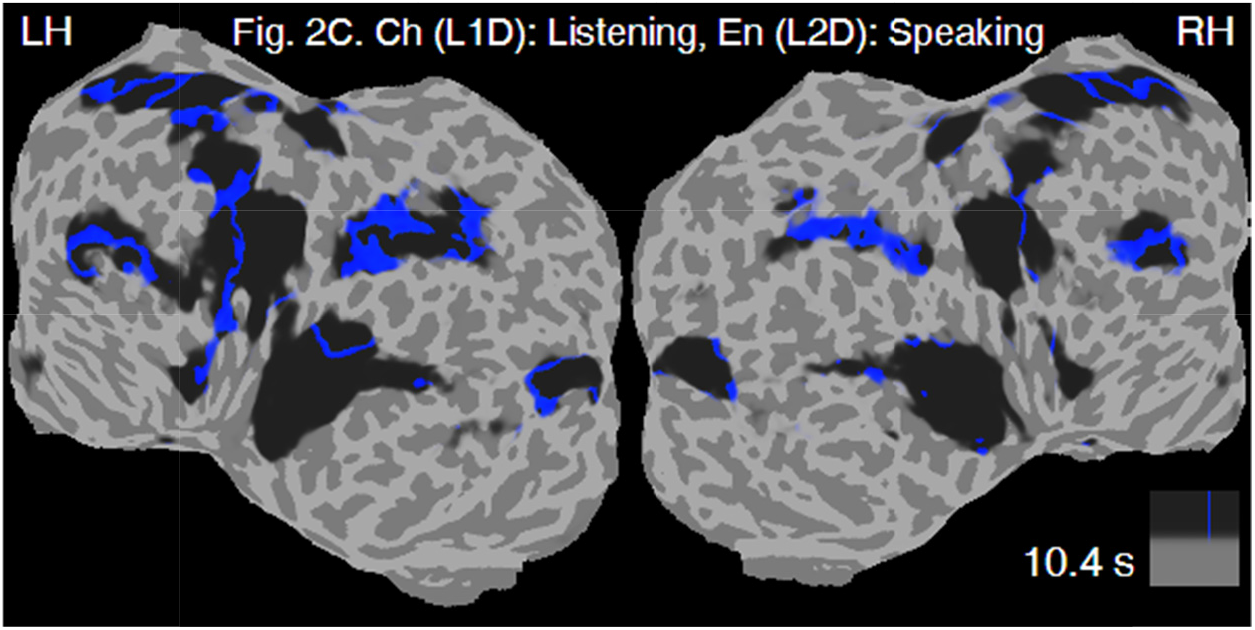
Brainstorms induced by an L1-to-L2 consecutive digit interpreting task.

**Movie S16.**
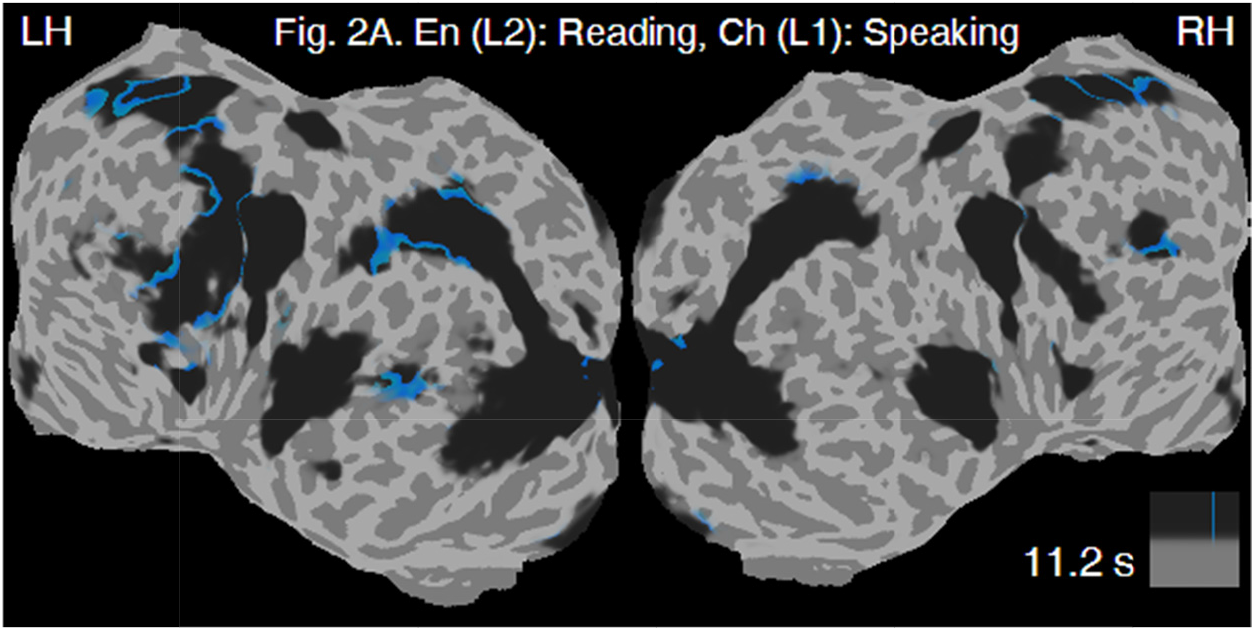
Brainstorms induced by an L2-to-L1 consecutive sight interpreting task.

**Movie S17.**
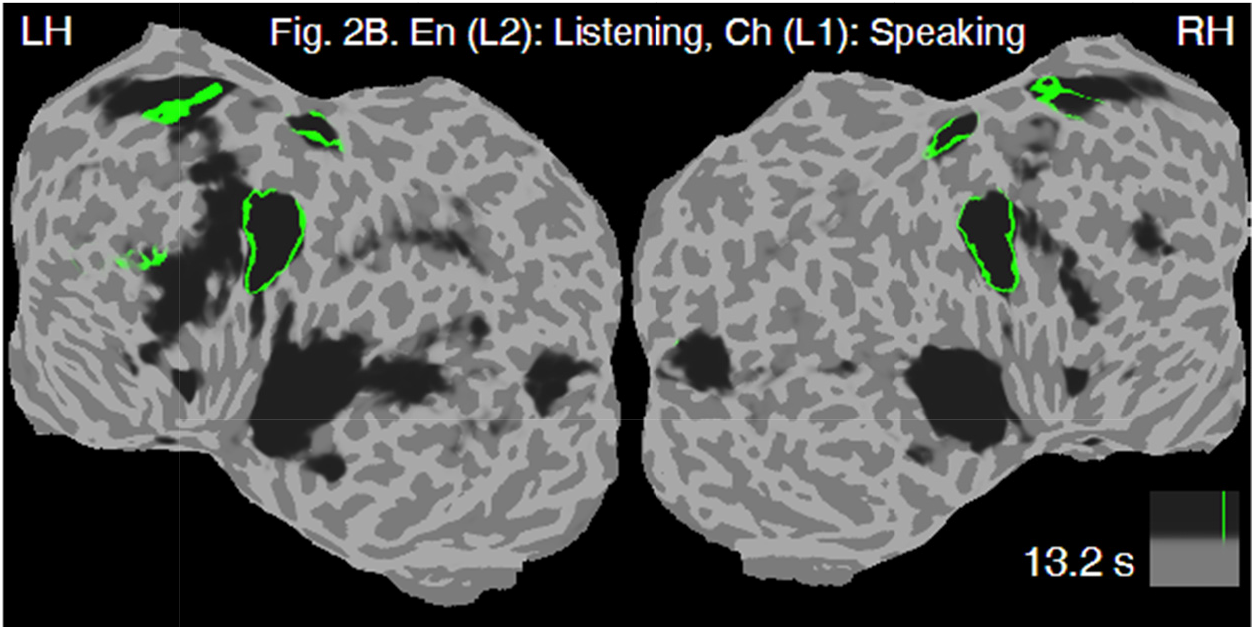
Brainstorms induced by an L2-to-L1 consecutive speech interpreting task.

**Movie S18.**
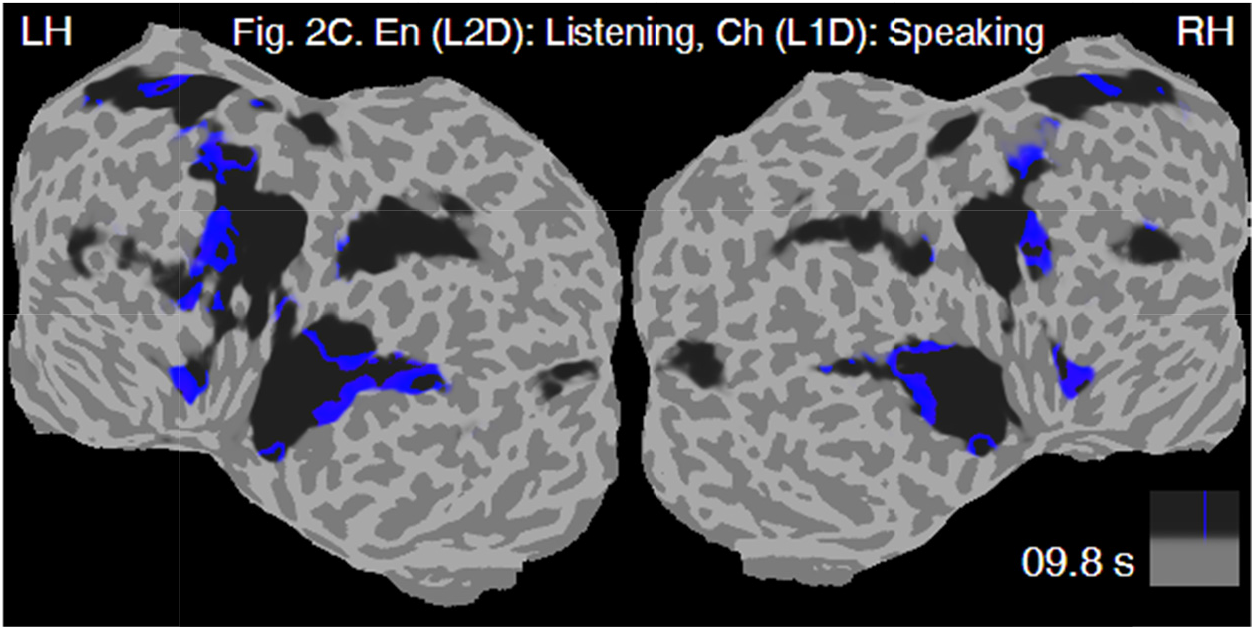
Brainstorms induced by an L2-to-L1 consecutive digit interpreting task.

**Movie S19.**
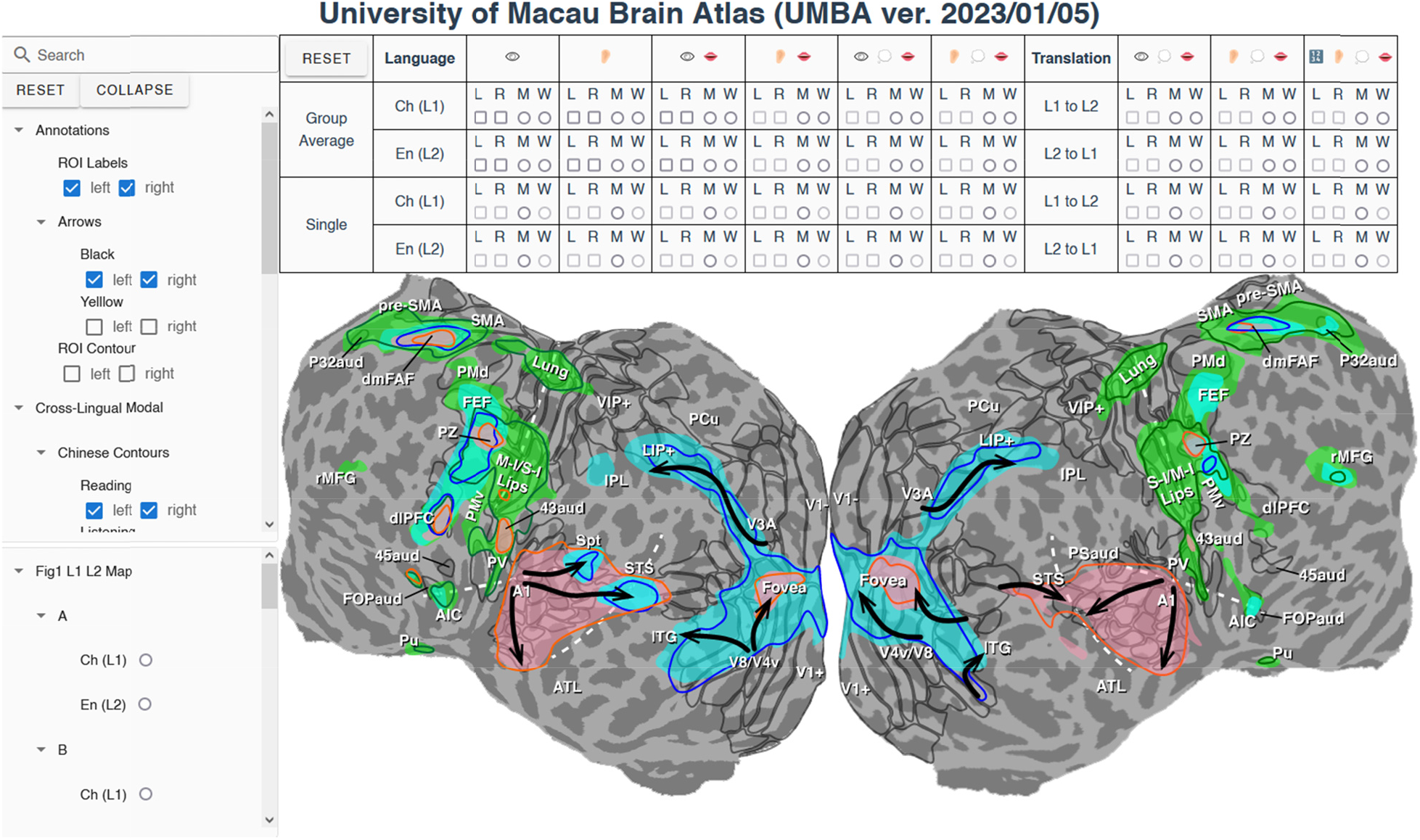
Demonstration of the University of Macau Brain Atlas (UMBA).

**Movie S20.**
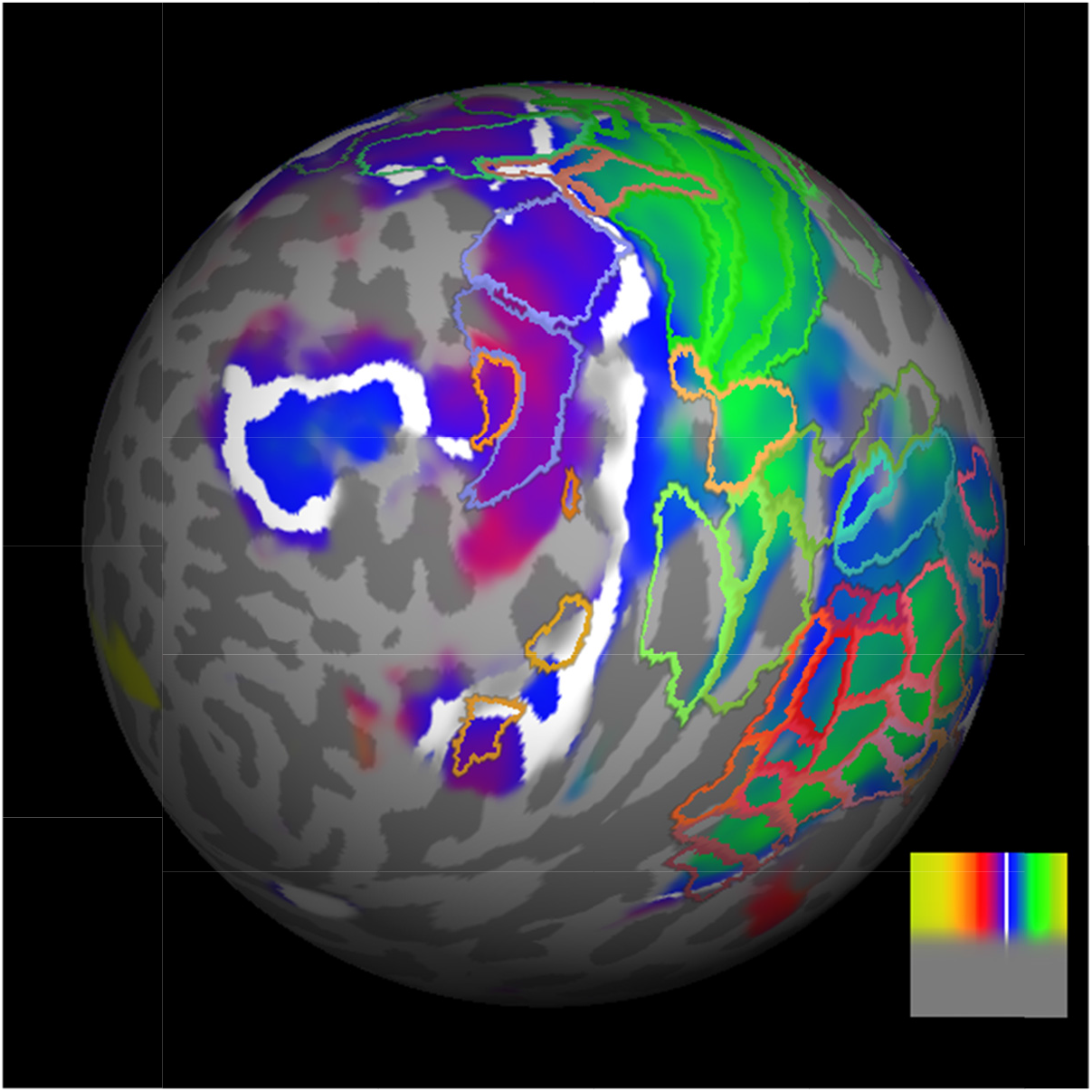
Brainsto ms on a spinning spherical cortical surface. The group-average activation map of the Chinese reading-memorizing-reciting task (see Fig. 1E) was displayed on the cortical surface of the left hemisphere of a representative subject. Color contours represent the borders of topological areas.

## Notes

### Competing Interest Statement

The authors have declared no competing interest.

